# Oncolytic virus-antibody combinations enhance immune-mediated killing of osteosarcoma

**DOI:** 10.64898/2026.09.10.750740

**Authors:** Tyler Barr, Lucia Cottone, Katherine Stanton, Victoria A. Jennings, Shaoli Das, Richard T. Baugh, Jonathan Stevenson, Hardev Pandha, Guy R. Simpson, Justas Stanislovas, Heather E. Owston, Dennis McGonagle, Peter V Giannoudis, Natasha J. Caplen, Sandra J. Strauss, Fiona Errington-Mais, Graham P. Cook

**Author notes:** These authors contributed equally to this work (Fiona Errington-Mais, Graham P. Cook).

## Abstract

A major barrier to effective immunotherapy in osteosarcoma (OS) is the highly immunosuppressive tumour microenvironment (TME), which limits immune recognition and elimination of tumour cells. We evaluated a panel of oncolytic herpes simplex viruses (oHSVs) for their direct oncolytic activity, immune-modulatory properties and capacity to counteract OS-associated immunosuppression using immunologically relevant *in vitro* models. We demonstrate that established OS cell lines, primary cell cultures and dissociated OS cells from freshly resected tumour samples are susceptible to direct oncolysis by three oHSVs; HSV1716, HSV1716-GMCSF, and HSV47Δ, although susceptibility levels varied. Treatment of peripheral blood mononuclear cells from healthy donors and OS patients with oHSVs enhanced natural killer (NK) cell activation and promoted immune-mediated killing of OS cell lines and primary OS cell cultures. Among the three viruses, HSV1716-GMCSF exhibited the strongest immune-stimulatory effects and was uniquely capable of reducing the abundance of CD163+CD206+ immunosuppressive TAMs; use of this oHSV was therefore prioritised. We developed a multicellular spheroid model of OS, incorporating OS cells, mesenchymal stem cells and TAMs, which exhibits resistance to immune-mediated killing, better reflecting the immunosuppressive TME in patients. In this model, pairing HSV1716-GMCSF treatment with either anti-GD2 or anti-EGFR monoclonal antibodies (mAbs), selected according to OS tumour antigen expression, significantly increased immune-mediated tumour cell killing. These findings suggest that personalised combination strategies pairing oHSVs with appropriate mAbs provide a promising therapeutic approach for OS by integrating direct oncolysis, remodelling of the immunosuppressive TME and enhanced immune-mediated tumour destruction.

## 1. Introduction

Osteosarcoma (OS) is the most common bone cancer in children and young adults, with peaks of incidence in adolescents and in those aged over 60 years (Mirabello et al., 2009; Stiller et al., 2013). First line treatments remain combinations of chemotherapy (high dose methotrexate, doxorubicin and cisplatin) both before and after surgical resection of the tumour (if appropriate based on the tumour site) (Strauss et al., 2021; Gerrand et al., 2025). The integration of systemic chemotherapy into standard of care treatment in the 1980s significantly improved patient outcomes. However, since then, patient survival rates have improved very little, with 5-year survival rates around 55% across all ages (Whelan et al., 2012). Patients diagnosed with metastatic disease, or those who go on to relapse following treatment have particularly poor outcomes, and 5-year survival is approximately 20-30% (Ferrari et al., 2003; Kager et al., 2003).

In 2011, the first new OS treatment for several decades, Mifamurtide (Mepact®), was approved as an adjuvant therapy in Europe, and is now available in Mexico, South Korea, Switzerland, and Israel (Ando et al., 2011; Anderson et al., 2014). Mifamurtide is a synthetic analogue of muramyl dipeptide (MDP), a component of Gram-positive bacterial cell walls encapsulated in phospholipid vesicles. Both MDP and Mifamutide are ligands of the intracellular pattern recognition receptor (PRR) NOD2 and trigger the production of pro-inflammatory cytokines and the induction of an immune response (Ando et al., 2011). Supporting the application of Mifamurtide (and indeed other immunotherapies) for OS treatment, there is mounting evidence for immune control of OS. Absolute lymphocyte counts post-chemotherapy are associated with improved survival of OS patients (Moore et al., 2010; Vasquez et al., 2017). Furthermore, specific immune cell infiltrates correlate with prognosis, for example higher levels of CD8+ T cells, M1-polarised macrophages and dendritic cells are associated with improved survival, suggesting that anti-tumour immunity can improve patient outcomes (Wang et al., 2026). By contrast, enrichment of myeloid-derived suppressor cells, regulatory T cells, and exhausted T cells are associated with an immunosuppressive tumour microenvironment (TME), disease progression and metastasis (Taylor et al., 2025).

Immune checkpoint molecules such as PD-L1 and CTLA-4 are expressed in OS, and their blockade can restore T-cell function and mediate tumour regression *in vitro* and *in vivo*. However, single agent immune checkpoint inhibitors have shown only modest benefit in patients (Lussier et al., 2015; Tawbi et al., 2017; Xie et al., 2020; Eghtedari et al., 2023; Han et al., 2025). By contrast, Mifamurtide therapy alongside chemotherapy has been demonstrated to improve 6-year overall survival from 70% to 78% (Meyers et al., 2008). This improved survival with Mifamurtide suggests that other immune-modulatory therapies have potential in OS, especially when used in combination with chemotherapy.

Oncolytic herpes simplex viruses (oHSVs) represent the most clinically advanced group of oncolytic viruses (OVs). These viruses exert anti-tumour activity through several complementary mechanisms including direct lysis of malignant cells and the activation of anti-tumour immune responses (Kaur et al., 2012; Workenhe et al., 2014; Andtbacka et al., 2016). One major advantage of oHSVs when compared to other OVs is their large DNA genome, which provides substantial capacity for genetic engineering. This allows the insertion of therapeutic transgenes designed to enhance immune stimulation and/or counteract immunosuppressive properties of the TME (Jennings et al., 2019; Wang et al., 2024; Tan et al., 2025).

To date, oHSVs have been approved for the treatment of melanoma and glioblastoma (O’Donoghue et al., 2016; Todo et al., 2022). Further development of oHSVs include insertion of alternative immune-modulatory transgenes encoding cytokines, antibodies and immune checkpoint inhibitors alongside novel delivery methods including nanoparticles and mesenchymal stem cell carriers (Zheng et al., 2025). Several preclinical studies have demonstrated the efficacy of multiple oHSVs against OS. However, these studies have exclusively utilised established human or murine OS cell line models of disease, either *in vitro* or *in vivo* (Bharatan et al., 2002; Currier et al., 2013; Wedekind et al., 2021; Ringwalt et al., 2024). The efficacy of oHSV in more representative human OS models has not been investigated, this represents a critical gap in our knowledge and in progress towards successful clinical translation.

We have utilised three distinct oHSVs; HSV1716, HSV1716-GMCSF and HSV47Δ. HSV1716 represents a first generation oHSV, with deletion of both copies of the *RL1* gene (encoding ICP34.5), reducing neurovirulent effects and improving cancer-selectivity (Valyi-Nagy et al., 1994). HSV1716-GMCSF retains the safety features of HSV1716 but is further engineered to deliver the transgene for GMCSF, this is designed to promote antigen presenting cell differentiation, such as dendritic cells. Finally, HSV47Δ, like HSV1716 has deletion of both copies of the *RL1* gene and an additional deletion of the α*47* gene. A clinically approved oHSV, similar to HSV47Δ(termed G47) is approved for the treatment of glioblastoma in Japan (Todo et al., 2022). We investigated the oncolytic and immune stimulatory activity of this panel of oHSVs in OS. Our experimental models included long established OS cell lines grown as both monolayers and spheroids, as well as primary patient derived cell cultures recently established from patient tumours and dissociated OS cells from freshly resected tumour tissue. In addition, OS cells were incorporated into more complex multicellular spheroids containing mesenchymal stem cells (MSCs) and tumour associated macrophages (TAMs) where they exhibited increased resistance to immune mediated killing. Importantly, using this more physiologically relevant microenvironment, we have identified rationally designed combination strategies pairing oHSV treatment with monoclonal antibodies (mAbs) targeting antigens overexpressed on OS cells. Critically, this novel combination approach enhances immune mediated destruction of OS. This represents a clinically deliverable combination strategy which warrants further investigation in clinical trials.

## 2. Materials and Methods

### 2.1 Cell lines, antibodies and reagents

All details of cell lines, primary cultures and patient samples including growth medium are provided in Supplementary Table 1. Details of other reagents including antibodies, fluorescent stains and buffers used in this study are detailed in Supplementary Table 2.

### 2.2 Cell culture

All cell cultures were routinely screened for mycoplasma using the LookOut® One Step Mycoplasma PCR Detection Kit (Sigma) and were consistently confirmed to be contamination free. Unless otherwise specified, growth media were supplemented with 10% heat inactivated foetal bovine serum (FBS; Sigma), with inactivation performed at 56 °C for 30 minutes prior to use.

### 2.3 Cell lines, primary cell cultures and patient samples

The OS cell lines MG-63 and HOS were studied alongside the primary OS cell cultures, SARC011, SARC050, SARC009 and SARC012, which were recently isolated from patient tumours. These samples were obtained from the UCL/UCLH Biobank for Studying Health and Disease, approved by the Health Research Authority (HRA) Research Ethics Service; Integrated Research Application System (IRAS) project identifier 272816, REC reference 20/YH/0088. This study was approved by the Biobank Ethical Review Committee (B-ERC) under project reference EC17.14. Samples were anonymised using Pro-Curo software (Pro-Curo Software Ltd., Horsham, UK). Matched primary patient tumour (OS-T-) and peripheral blood (OS-B-) samples were obtained from the Royal Orthopaedic Hospital Research Tissue Bank; ethical approval reference 22/EM/0042 issued by East Midlands/Derby Research Ethics Committee. Healthy donor leukocyte apheresis cones were supplied by the National Health Service Blood and Transplant unit (NHSBT). PBMCs were isolated using Lympholyte^TM^ (Cedarlane) and density gradient centrifugation.

Bone marrow-derived mesenchymal stem cells (MSCs) were isolated from human healthy donor bone marrow aspirates. Ethical approval was obtained from the NREC Yorkshire and Humberside National Research Ethics Committee (18/YH/0166). Isolation and analysis of MSCs was as previously described (Wilson et al., 2024).

### 2.4 Dissociation of OS tumour tissue and establishment of primary cell cultures

Patient OS tumour tissue (collected during surgery) was transported in University of Wisconsin (UW) solution and kept at 4°C until dissociation: tissue was washed twice in PBS, macerated using a scalpel and dissociated for 45–60 minutes using 5 mg/mL collagenase II (Invitrogen) primary cell culture growth medium without FBS at 37°C. Cells were then washed, a single cell suspension obtained using a cell strainer and cells cultured in complete growth medium (Supplementary Table 1). For establishment of primary cell cultures, SARC009, SARC011, SARC012 and SARC050, cells were maintained in complete growth medium and routinely passaged upon near confluence. OS primary cells were confirmed to carry genomic aberrations using whole exome sequencing or shallow whole genome sequencing.

### 2.5 Spheroid cultures

Monoculture spheroids were generated by seeding 1 x 10^4^ OS cells into a 96 well U bottom ultra-low adhesion plate (Corning Incorporated) for 7 days. Multicellular spheroids were generated by combining 1 x 10^4^ OS cells with 5 x 10^3^ CD14+ monocytes and 1 x 10^3^ MSCs for 7 days.

### 2.6 *In vitro* generated TAMs

TAMs were generated by co-culture of healthy donor PBMCs with HOS or MG-63 OS cell lines at a ratio of 50:1 for 7 days, as previously described (Jennings et al., 2024). CD14+ cells were then isolated using CD14 magnetic beads (Miltenyi Biotec), and the purity of selection validated by staining with anti-CD14 antibody and flow cytometry.

### 2.7 Lentiviral transduction of cell lines and primary cell cultures

Lentivirus was generated using GLV3-CMV-[ORF]-PGK-Puro vector (GenScript). Firefly luciferase (Luc2)*-*expressing MG-63, HOS, SARC011 and SARC050 cells were generated by lentiviral transduction. Briefly, cells were seeded into 100mm dishes over night to reach ∼80% confluency. Lentivirus was then added to cells at an MOI of 10 with 8 µg/mL polybrene in serum-free Dulbecco’s Modified Eagles Medium overnight. The following day, media was removed from wells and replaced with complete growth medium. Cells were selected in puromycin at the concentrations indicated in Supplementary Table 1.

### 2.8 Oncolytic viruses

HSV1716 and HSV1716-GMCSF were obtained from Virttu Biologics. HSV47Δ was made by GRS and HP at the University of Surrey. HSV1716, HSV1716-GMCSF and HSV47Δ were amplified on Vero cells. A standard plaque assay on the Vero cell line was used to assess the titre of virus stocks and HSV-treated cell lysates. Reovirus type 3 Dearing strain (Reolysin) stocks were provided by Oncolytics Biotech and virus was titred by plaque assay on L929 cells. All virus stocks were stored at -80°C until use.

### 2.9 Assessment of cell viability and growth inhibition

To quantify remaining adherent viable cells after treatment of monolayer cultures with oHSVs, supernatants were removed, cells were washed with phosphate buffered saline (PBS) and fixed in 1% paraformaldehyde (PFA) in PBS. Cells were then stained with 1% methylene blue in ethanol:water (1:1) for 3 minutes and washed in water. Plates were air dried and imaged using BIO-RAD ChemiDoc^TM^ MP imaging system. ImageJ V.1.52a software was used to analyse the inverted image of plates, where integrated density of untreated and treated wells was determined. Background integrated density was determined using empty wells and subtracted from all values. The integrated density of treated cells was expressed as a percentage relative to signal from untreated wells. CellTiter-Glo® reagent (Promega) was used to determine growth inhibition of OS spheroid cultures and dissociated OS cells from primary patient samples, as per manufacturers’ instructions.

### 2.10 Flow cytometry-based phenotyping

For the assessment of NK cell activation, after PBMC treatments, cells were stained with anti-CD3, anti-CD56, ant-CD69 and anti-CD317 antibodies. For the assessment of antigen expression on OS cells, cells were stained with anti-EGFR, anti-HER2, anti-GD2 and anti- CD20 antibodies. For the assessment of TAM phenotype, cells were stained with anti-CD14, anti-CD206 and anti-CD163 and anti-HLA-DR (MHC class II). All antibodies were added to cells for 30 min at 4 °C in FACs buffer, then washed in PBS and fixed in 1% PFA. Details of all antibodies/ buffers in Supplementary Table 2. For all flow cytometry-based assays, cells were analysed using a Cytoflex LX (Beckman Coulter; Brea, CA, USA) flow cytometer and analysed using FlowJo V10.8.1 and CytExpert V2.5 software.

### 2.11 NK cell degranulation assays

PBMC were co-cultured with OS tumour cell targets with or without TAMs at a ratio of 10:1:1 for 1 hour, before the addition of anti-CD3, anti-CD56 and anti-CD107a antibodies and Brefeldin A at a final concentration of 3 µg/mL (Supplementary Table 2). Co-cultures were incubated for a further 3 hours, and cells were then washed in PBS, fixed in 1% PFA, and analysed using flow cytometry.

### 2.12 Immune killing assays

For functional immune killing assays, healthy donor PBMC were activated with OV or Mifamurtide, at concentrations or multiplicity of infection (MOI) specified for individual experiments, for 48 hours. For luciferase-based killing assays, PBMC were then co-cultured with Luc2-expressing cell lines or primary cell cultures at a ratio of 20:1 for 24 hours. D-luciferin was added to wells at 150 µg/mL and luminescence measured using a Cytation 5 plate reader. Relative luminescence was calculated by subtracting background signal (wells containing media and d-luciferin only) from all samples, then calculating relative luminescence of treated wells as a percentage of untreated wells. For flow cytometry based immune killing assays, PBMC were co-cultured with CellTracker Green CMFDA-stained OS target cells at a ratio of 20:1 for 24 hours, and then stained with LIVE/DEAD™ Fixable Yellow Dead Cell Stain Kit, and the percentage of dead tumour cells quantified by flow cytometry, as previously described (Barr et al., 2025).

### 2.13 Enzyme-linked immunosorbent assays (ELISA)

IFNα, IL-6 and GM-CSF ELISAs was carried out using Maxisorp plates and matched paired antibodies (Supplementary Table 2). Detection of IL-1β in cell supernatants was carried out using Human IL-1 beta/IL-1F2 Quantikine ELISA Kit (Bio-techne).

### 2.14 Transcriptome Analysis

Bulk mRNA sequencing of OS in vitro generated TAMs was performed on *in vitro* generated TAMs produced by co-culture with HOS and MG-63 cell lines. Total RNA was isolated from CD14-selected TAMs using RNeasy Mini Kit (Qiagen), and RNA quantification was performed using Qubit Fluorometry HS kit (Thermo Fisher Scientific). mRNA sequencing was performed by Novogene using Illumina Sequencing PE150 of 400 ng of total RNA per sample. Raw reads were aligned to the *Homo sapiens* reference genome (GRCh38/hg38) using HISAT2 v2.0.5 with default parameters. Gene-level read counts were obtained using featureCounts v1.5.0-p3, and transcript abundance was normalised as fragments per kilobase of transcript per million mapped reads (FPKM).

For scRNAseq analysis, osteosarcoma snRNA-seq/scRNA-seq data were downloaded from the Alex’s Lemonade Stand Foundation (ALSF) Single-cell Pediatric Cancer Atlas (ScPCA) (Hawkins et al., 2026). The dataset SCPCP000023 comprised snRNA-seq data from 30 primary patient tumour samples. The snRNA-seq data were processed in Seurat v5, following the preprocessing and integration steps of the CCBR SINCLAIR pipeline (https://zenodo.org/records/20146239). The following parameters were used for initial QC filtering: minimum number of genes with at least one molecule detected in a cell >200, minimum number of UMIs detected in a cell >700, and maximum percentage of mitochondrial transcripts in a cell <30%. Samples with <50 cells passing these QC criteria were removed from the analysis. A total of 16 samples passed these QC criteria and were processed for downstream analysis. Samples were integrated in Seurat using the Harmony algorithm, and the integrated data were SCTransformed, followed by PCA, clustering, and UMAP. A clustering resolution of 0.2 was chosen as the optimal resolution based on average silhouette width calculation. Per-cell gene set enrichment for immune cell types was calculated using UCell (Andreatta et al., 2021) with cell-type signatures from the CellMarker database (Hu et al., 2023).

Bulk RNAseq data from patient tumours was obtained from the TARGET-OS dataset downloaded from the NCI Genomic Data Commons (GDC) (Heath et al., 2021). The dataset used in this study is available through the database of Genotypes and Phenotypes (dbGaP) under the accession number phs000218. Gene expression (UQ-FPKM) data was downloaded for the selected candidate target genes (EGFR, ERBB2, B4GALNT1).

### 2.15 Statistical analysis

Significant differences in results were determined using a Student’s t-test, one-way ANOVA or two-way ANOVA. Statistical analyses were performed using GraphPad PRISM 10 software.

## 3. Results

### 3.1 Oncolytic HSVs exhibit direct oncolytic effects against OS cell lines and primary cell cultures

We investigated the oncolytic activity of three distinct herpes simplex virus-based oncolytic viruses (oHSVs); HSV1716, HSV1716-GMCSF and HSV47Δ against two long-established OS cell lines (MG-63 and HOS) and four primary OS cell cultures more recently established from patient tumours (SARC009, SARC011, SARC012 and SARC050). These primary OS cell cultures were derived from tumour samples representing various OS-subtypes and both chemotherapy-treated or treatment-naïve tumours (Supplementary Table 1).

Monolayer cultured cells were treated with oHSVs at 0.1 or 1 plaque forming units (PFU)/cell for up to seven days and methylene blue staining was used to quantify the remaining cells relative to untreated controls. Treatment with all three oHSVs at 1 PFU/cell for 72 hours and seven days significantly decreased the proportion of remaining viable cells when compared to untreated controls (Figure 1A-B). Similar results were also observed following treatment with a lower MOI of 0.1 PFU/cell (Supplementary Figure 1). Different levels of oncolysis were observed between the cell lines and between the three oHSVs, consistent with genetic differences in both cells and viruses. Compared to results at seven days, oncolysis was lower at 72 hours or with low level initial infection (0.1 PFU/cell), consistent with a need for viral replication and reinfection with progeny virions for maximum effect. Productive infection of all three oHSVs was confirmed for each of the six OS cell cultures using plaque assay of lysates on Vero cells (Supplementary Figure 1D). Clear cytopathic effects were observed (rounding and detachment) in response to HSV1716 and HSV1716-GMCSF infection (Figure 1C). By contrast, HSV47Δ treatment of primary cell cultures resulted in syncytia formation (Figure 1C). Furthermore, the two GFP-encoding oHSVs (HSV1716 and HSV47Δ), confirmed expression from the viral genome (Figure 1C).

**Figure 1:**
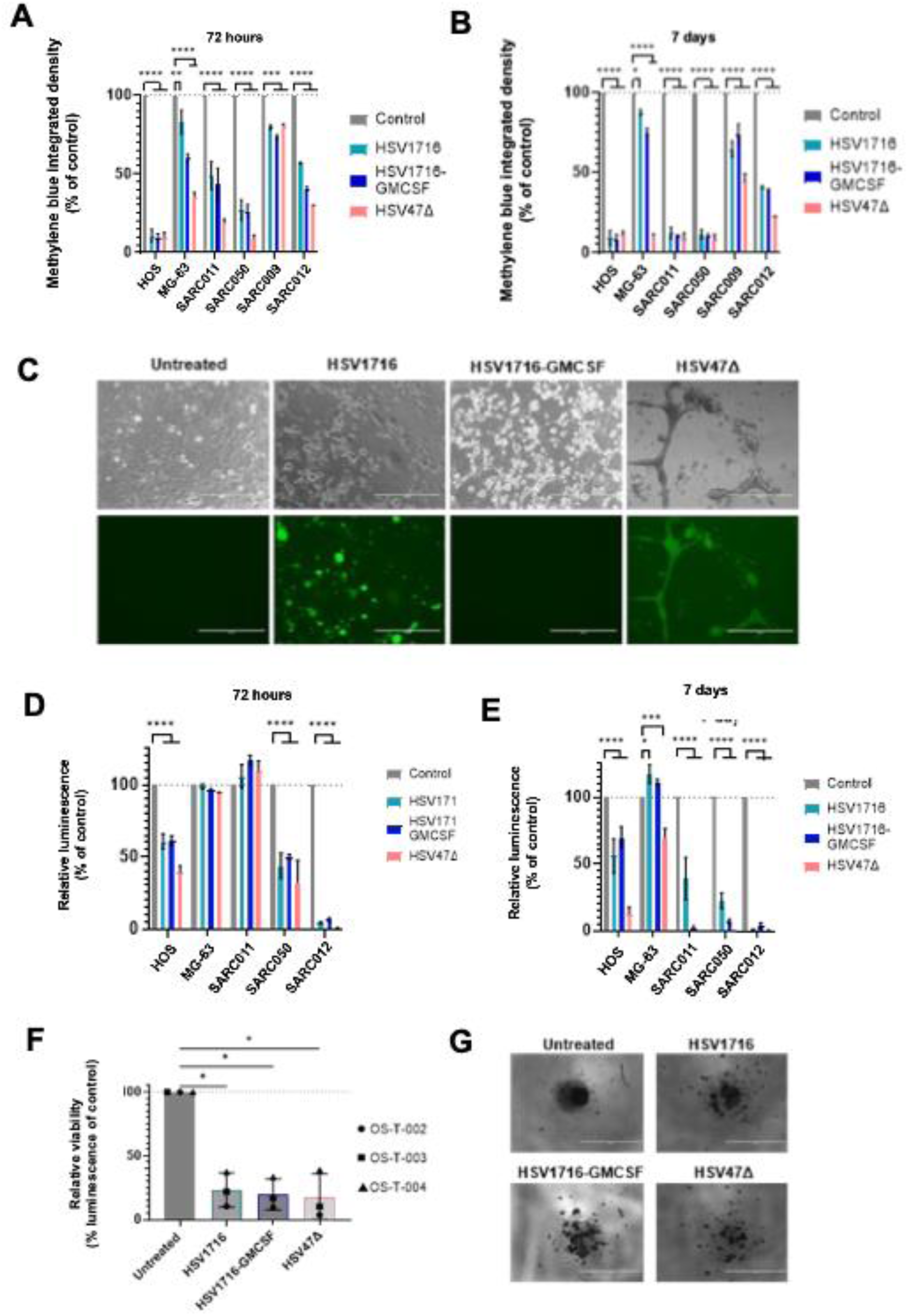
oHSVs exhibit direct oncolytic effects against OS cell lines and primary cell cultures. (A-B) MG-63 and HOS OS cell lines and SARC011, SARC050, SARC009 and SARC012 primary cell cultures were treated ± HSV1716-GFP, HSV1716-GMCSF or HSV47Δ at 1 PFU/cell for (A) 72 hours or (B) 7 days. Monolayers were then stained with methylene blue, and the proportion of remaining viable cells was quantified by using ImageJ software to calculate integrated density of staining relative to untreated controls, mean ± SEM for n=3 independent experiments, analysed using two-way ANOVA. (C) SARC050 cells were treated ± oHSVs at 1 PFU/cell for 72 hours. Cells were then imaged using EVOS microscope at 10x magnification, scale bar = 400 µm. (D-E) MG-63, HOS, SARC011, SARC050 and SARC012 were seeded into low attachment 96 well plates for 7 days to generate spheroid cultures. (F-G) OS patient tumours (OS-T-002, OS-T-003 and OS-T004) were digested in collagenase II and single cell suspensions were then added to low attachment 96 well plates. (D-G) Cells were treated ± HSV1716-GFP, HSV1716-GMCSF or HSV47Δ at 1 PFU/cell for (D) 72 hours or (E-G) 7 days. (D-F) Cell Titre-Glo® reagent was then added to wells and luminescence measured using a plate reader. Results shown as percent luminescence relative to untreated controls. (G) OS-T-002 was imaged using EVOS microscope at 4x magnification, scale bar = 1 mm. (D-E) mean ± SEM for n=3 independent experiments, analysed using two-way ANOVA.

Whilst 2-dimensional (2D) *in vitro* models offer an effective tool for screening of novel agents, 3D models are more representative of patient tumours. For example 3D tumour architecture impedes drug penetration and dissemination throughout the structure, leading to increased drug resistance (Munoz-Garcia et al., 2021). We tested the oncolytic activity of oHSVs in OS cells cultured as 3D spheroids. Spheroids were treated with oHSVs at 1 PFU/cell for up to 7 days, and the effect on cell viability was measured using Cell Titre-Glo. Three spheroid cultures (HOS, SARC012 and SARC050) demonstrated susceptibility to all three oHSVs after 72 hrs (Figure 1D). In contrast, the MG-63 cell line and SARC011 primary OS culture were resistant to direct oncolytic activity of all three oHSVs after 72 hrs (Figure 1D). However, by 7 days, SARC011 cells showed significantly increased cell death following treatment with all three oHSV, whilst the MG-63 cell line remained resistant to HSV1716 and HSV1716-GMCSF (Figure 1D-E). These varied responses highlight the importance of testing agents across multiple samples in more representative culture conditions. We validated these findings in OS tumour cells isolated directly from patients less than 48 hours prior to tissue dissociation and treatment with oHSVs. Tumours were obtained from consenting patients who were undergoing routine surgical resection of their tumour. Importantly, these patients had recently (≤28 days prior to surgery) undergone initial rounds of neoadjuvant methotrexate/doxorubicin/cisplatin (MAP) chemotherapy, meaning that any remaining viable cells used in these assays would represent a population of treatment-refractory tumour cells with the potential for relapse.

Tumour samples from three patients were digested in collagenase II and single cell suspensions were seeded into low adhesion plates and treated with oHSVs for up to 7 days; treatment with oHSVs significantly reduced cell viability in all three patient-derived samples after 7 days of treatment (Figure 1F and Supplementary Figure 1E). An untreated sample (OS-T-002) showed formation of spheroid-like architecture after seven days of culture; this spheroid formation was inhibited by oHSV treatment and instead cells appear dispersed and disaggregated (Figure 1G). Our results, using a range of *in vitro* model systems (established cell lines, recently established primary cell cultures and fresh tumour tissue) show that OS cells are highly sensitive to the direct oncolytic activity of oHSVs.

### 3.2 Oncolytic HSVs stimulate immune activation and immune-mediated killing of OS

The direct oncolytic activity of oHSVs only represents one of their multimodal mechanisms of action. Importantly, oHSVs have been widely demonstrated to activate immune-mediated killing of cancer cells in both *in vitro* and *in vivo* models as well as in patients (Workenhe et al., 2014; Andtbacka et al., 2016; Jennings et al., 2019; Tan et al., 2025). The approval of Mifamurtide for the treatment of OS in 2011 provides scope and impetus for the use of additional immune-modulatory agents for OS.

To compare the immune stimulatory properties of each oHSV to Mifamurtide, PBMC from healthy donors were treated with oHSV or Mifamurtide for 48 hours, and the percentage of activated NK cells (as determined by expression of cell surface CD69) was quantified using flow cytometry. Each of the oHSV vectors significantly increased NK cell activation when compared to the untreated control and to Mifamurtide treatment (Figure 2A). Although much less potent than oHSVs, Mifamurtide treatment still resulted in significant NK cell activation when compared to the untreated control (p<0.05; Figure 2A).

**Figure 2:**
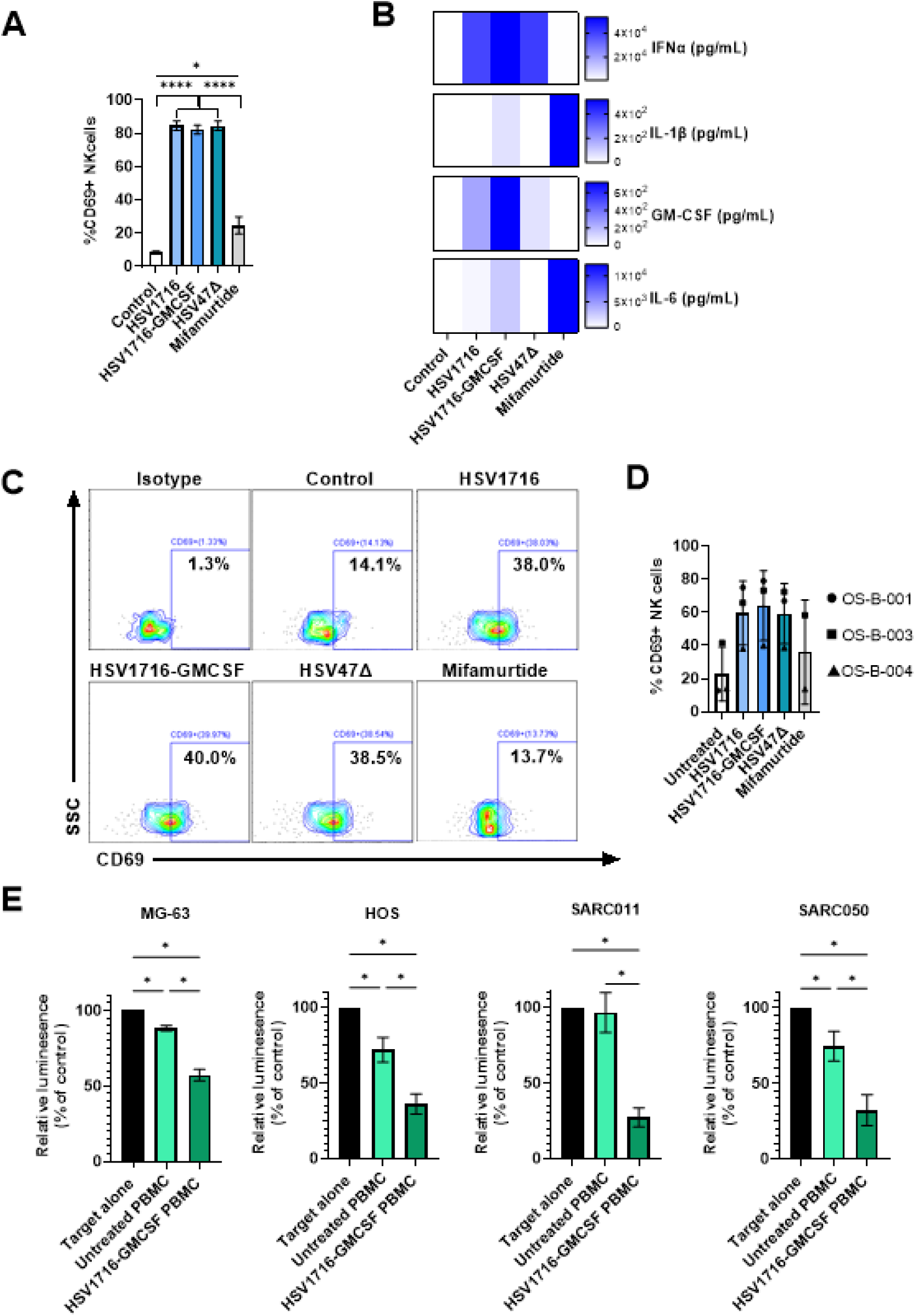
oHSVs induce immune activation and immune-mediated killing of OS. (A-B) PBMCs from healthy donors were treated with HSV1716-GFP, HSV1716-GMCSF, HSV47Δ at 0.1 PFU/cell or Mifamurtide at 100 µM for 48 hours. (A) PBMC were stained with anti-CD3, anti-CD56 and anti-CD69 antibodies, and the percentage of CD56+ CD3-CD69+ activated NK cells was determined by flow cytometry, presented as mean ± SEM for n=4 PBMC donors, analysed using one-way ANOVA. (B) PBMC supernatants were collected and screened for IFNα, GM-CSF, IL-1β and IL-6 using ELISA; heat maps represent mean concentration from n=4 independent PBMC donors. (C-D) PBMCs from OS patients were treated with HSV1716-GFP, HSV1716-GMCSF, HSV47Δ at 0.1 PFU/cell or Mifamurtide at 100 µM for 48 hours. PBMC were stained with anti-CD3, anti-CD56 and anti-CD69 antibodies, and the percentage of CD56+CD3-CD69+ NK cells was determined by flow cytometry. (C) Representative flow cytometry gating for OS-B-004 and (D) results presented as mean ± SEM, where symbols represent individual samples. (E) Healthy donor PBMC were activated ± HSV1716-GMCSF at 0.1 PFU/cell for 48 hours, PBMC were then co-cultured with Luc2-expressing MG-63, HOS, SARC011 or SARC050 cells for 24 hours at a ratio of 20:1. D-luciferin was added to plates at 150 µg/mL and luminescence measured using a plate reader. Results presented as luminescence relative to none co-cultured target cell controls, mean ± SEM, n≥4. Statistical comparison performed using one-way ANOVA.

To explore cytokine secretion, cell-free supernatants from oHSVs or Mifamurtide-treated PBMC were screened for IFNα, GM-CSF, IL-1β and IL-6 using ELISA. oHSV treatment was associated with high levels of IFNα secretion, whereas Mifamurtide induced both IL-1β and IL-6, as expected (Figure 2B). Notably, HSV1716-GMCSF treatment resulted in high levels of GM-CSF, IL-1β and IL-6 (Figure 2B). While GM-CSF is not directly associated with IL-1β and IL-6 production, GM CSF expands and activates myeloid cells, and can thus amplify their response to viral sensing, this could explain the more diverse cytokine response induced by this virus compared with HSV1716 and HSV47Δ (Bouzeineddine et al., 2025). These data demonstrate that HSV1716, HSV47Δ and Mifamurtide treatment result in distinct cytokine signatures. However, HSV1716-GMCSF may offer a treatment strategy which promotes both the immune-stimulatory type-I IFN response associated with OV therapy as well as the macrophage activating/ polarising cytokine signature associated with Mifamurtide treatment.

A key concern for immunotherapy-based treatment of OS is the effect of first line chemotherapy treatments (MAP) on lymphocyte counts in patients and the impact on efficacy of immune-stimulation in patients following immunotherapy. Therefore, we assessed the ability of oHSV to activate PBMCs isolated from three OS patients who had completed MAP chemotherapy ≤1 month prior to the blood sample collection; oHSVs activated patient-derived NK cells from all three samples, although these results did not reach statistical significance. We obtained sufficient PBMC from two of three patients to perform an additional stimulation with Mifamurtide treatment which also induced NK cell activation; however, the mean level of CD69 expression was lower than that seen with oHSVs (Figure 2C-D).

Together, these data suggest that prior chemotherapy treatment does not substantially hinder NK cell activation by oHSVs in patients, further validating the potential applicability of oHSVs in OS. In addition, oHSVs showed greater NK cell activation capacity than Mifamurtide. However, the strong myeloid-activating capacity of Mifamurtide remains attractive in myeloid-rich tumours such as OS. Accordingly, HSV1716-GMCSF may provide a dual approach to exploit both axes, simultaneously enhancing NK-cell activation whilst also modulating macrophage populations. This OV was therefore prioritised in further investigations. Finally, we tested the ability of HSV1716-GMCSF activated PBMCs to stimulate immune-mediated killing of firefly luciferase (*Luc2*)-expressing OS tumour cell targets. These data demonstrated that HSV1716-GMCSF treatment of healthy donor PBMCs significantly increased immune-mediated killing of OS cell lines (MG-63 and HOS) and primary cell cultures (SARC011 and SARC050) (Figure 2E). Consistent with an enhanced capacity to activate NK cells, HSV1716-GMCSF was superior to Mifamurtide in stimulating immune-mediated killing of SARC011 and SARC050 primary OS cell cultures (Supplementary Figure 2).

### 3.3 *In vitro* generated OS TAMs are repolarised by HSV1716-GMCSF treatment

Oncolytic viruses, especially those encoding particular transgenes, offer enhanced immune stimulatory/ modulatory effects on cytotoxic immune cells whilst simultaneously repolarising macrophage populations (Jennings et al., 2019; Cao et al., 2021; Wang et al., 2023; Jennings et al., 2024). We sought to explore the direct impact of HSV1716-GMCSF on macrophage populations using a recently developed *in vitro* model of OS generated TAMs, produced following co-culture of HOS or MG-63 cell lines with healthy donor PBMC. After 7 days, CD14+ cells were isolated by magnetic selection and cells were stained for CD14, CD163, CD206 and MHC class II to quantify the proportion of cells with a TAM-like phenotype. Co-culture of PBMCs with HOS and MG-63 OS cell lines resulted in a significant increase in CD14+CD206+CD163+ macrophages (Figure 3A-B). Moreover, the generated macrophages had significantly reduced MHCII expression when compared with controls (Figure 3A-B); we denote these *in vitro* generated TAMs as HOS-TAM and MG-63-TAM respectively.

**Figure 3:**
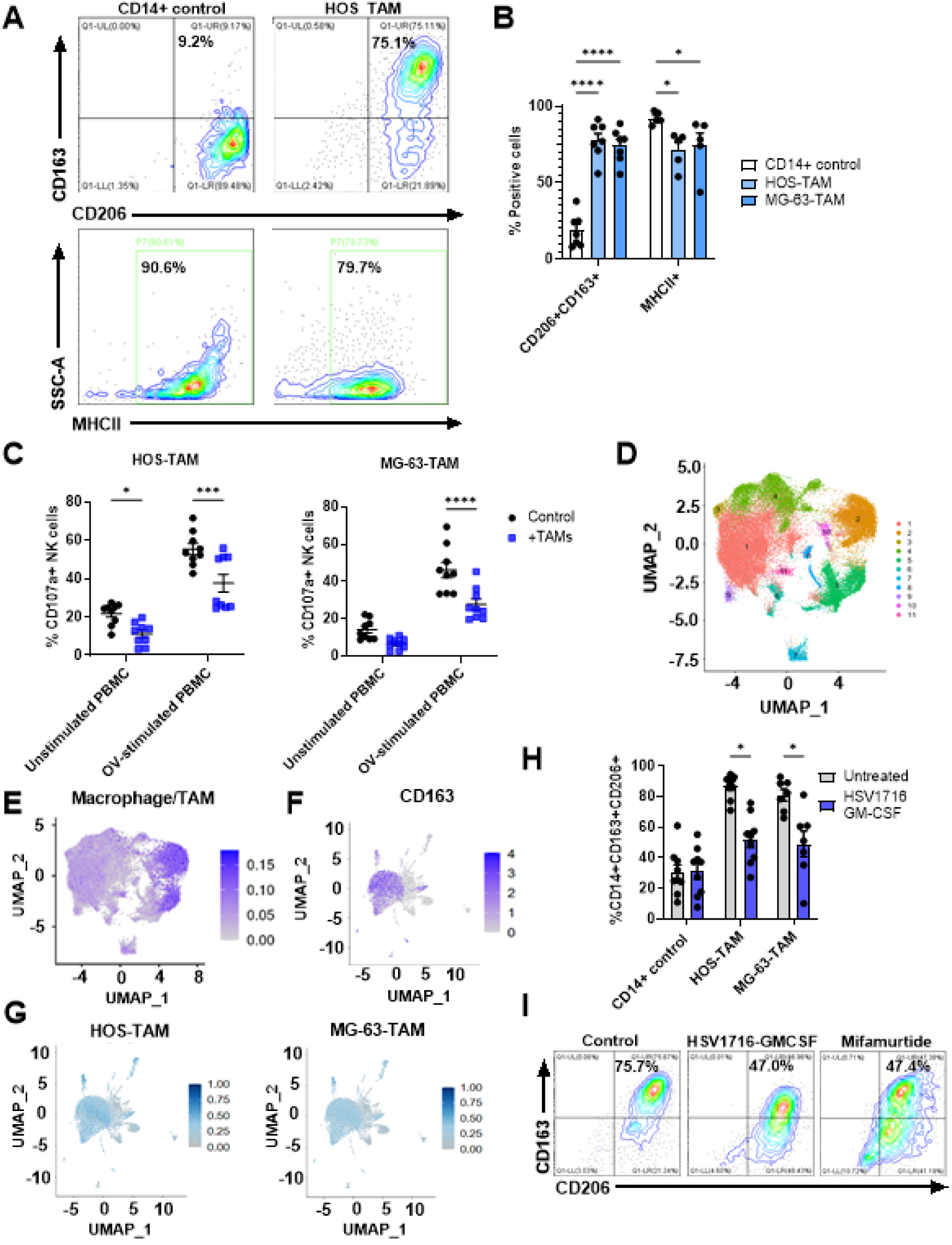
HSV1716-GMCSF repolarises *in vitro* generated OS TAMs. (A-C) OS TAMs were generated by co-culture of HOS or MG-63 OS cell lines with healthy donor PBMCs at a ratio of 1:50 for 7 days. CD14+ cells were isolated using magnetic bead selection. (A-B) Cells were stained for CD14, CD163, CD206 and MHC class II (MHCII) using fluorescently conjugated antibodies and assessed by flow cytometry. (A) Representative flow cytometry plots for CD14+ cells alone (not co-cultured with HOS) control and HOS-TAMs. (B) Summary data for cell phenotypes from n≥5 independent PBMC donors represented by symbols, bars show mean ± SEM, statistical comparison performed using two-way ANOVA. (C) HOS or MG-63 target cells were cultured with unstimulated or reovirus-stimulated PBMCs, ± HOS-TAM or MG-63-TAM (1:10:1) for 4 hours. Cells were stained with anti-CD3, anti-CD56 and anti-CD107a antibodies, and the percentage of degranulating NK cells quantified by flow cytometry, n=9 PBMC donors represented by symbols, bars show mean ± SEM, statistically analysed using two-way ANOVA. (D) Single cell RNA sequencing data from 16 OS patient tumours presented in a UMAP showing 11 distinct clusters. (E) Clusters 2 and 5 were identified as macrophage/TAM populations and (F) subclustered to identify CD163+ population of cells. (G) Bulk RNA sequencing data from HOS-TAM and MG-63-TAM were analysed and top 100 most highly expressed genes identified, enrichment of these genes is shown in OS scRNAseq data macrophage/TAM subcluster. (H-I) Day 7 HOS-TAM and MG-63-TAM were isolated using CD14+ magnetic selection and co-cultured with HOS or MG-63 cell line at 1:1. Cells were treated ± HSV1716-GMCSF at 0.1 PFU/cell for 48 hours and stained for CD14, CD163 and CD206 using fluorescently conjugated antibodies and assessed by flow cytometry, n=9 PBMC donors represented by symbols, bars show mean ± SEM, statistically analysed using two-way ANOVA. (I) Representative flow cytometry plot for HOS-TAM treated ± HSV1716-GMCSF at 0.1 PFU/cell or Mifamurtide at 100 µM.

An important role of TAMs is suppression of cytotoxic immune cell populations. To validate the functional immunosuppressive nature of our *in vitro* generated TAMs, CD14+ isolated TAMs were co-cultured with PBMC and OS cell lines and NK cell degranulation was assessed as a measure of NK cell function; both HOS-TAM and MG-63-TAM were able to inhibit NK cell degranulation, confirming their immunosuppressive capacity (Figure 3C). We then compared the HOS-TAM and MG-63-TAM cells to TAMs present in OS samples using publicly available single cell RNA sequencing (scRNAseq) data. UMAP analysis identified 11 distinct clusters within data from 16 OS primary tumour samples and macrophage/TAM populations were identified as clusters 2 and 5 (Figure 3D and E), with expression of CD163 confirmed within these clusters when analysed in isolation (Figure 3F). We then performed bulk RNA sequencing on HOS-TAM and MG-63-TAM populations and identified the top 100 most highly expressed genes in these cells. This gene signature was then tested in the OS scRNA-seq data. Both the HOS-TAM and MG-63-TAM gene signatures were enriched in primary OS TAMs (Figure 3G), demonstrating that the *in vitro* generated TAMs were representative of the OS patient TAM populations, confirming the relevance of the *in vitro* TAM model used in this study.

Having validated the *in vitro* OS TAM model, we next assessed the effect of HSV1716- GMCSF treatment on TAM polarisation (Figure 3H). To do this, HOS-TAM and MG-63-TAM were isolated using CD14+ selection and then re-cultured with HOS or MG-63 cell lines respectively, at a ratio of 1:1. Cells were then treated with HSV1716-GMCSF at 0.1 PFU/cell for 48 hours and TAM phenotype was examined. Treatment of TAMs with HSV1716-GMCSF significantly reduced the abundance of CD14+CD206+CD163+ TAMs when compared with untreated controls (Figure 3H). Furthermore, the TAM repolarisation observed with HSV1716-GMCSF was similar to that observed with Mifamurtide treatment (Figure 3I).

### 3.4 Multicellular spheroid models of OS exhibit resistance to immune-mediated killing

Next, we aimed to incorporate our TAM model into a preclinical system which better mimics the immunosuppressive nature of the OS TME. The development of a more refined OS models would allow us to test rationally designed combination strategies, alongside HSV1716-GMCSF or other OV. As such, a multicellular spheroid model was developed, incorporating TAMs, and mesenchymal stem cells (also known as mesenchymal stromal cells; MSCs), to mimic the supportive stromal niche that modulates osteosarcoma proliferation and treatment-resistance (Xu et al., 2009; Tu et al., 2012). To investigate immune mediate killing, we selected a model system that would allow us to distinguish between immune-mediated killing and oncolysis. As both HOS and MG-63 cell lines are susceptible to oncolysis by HSV-based OV but MG-63 cells are resistant to reovirus oncolysis (Supplementary Figure 3), reovirus was used to activate immune cells for initial characterisation of the model. Firefly luciferase-expressing MG-63 cells were cultured with CD14+ monocytes (to generate TAMs) and MSCs at a ratio of 10:5:1 for 7 days in low adhesion plates to generate spheroid structures (Figure 4A). Multicellular (MG- 63+TAM+MSC) or MG-63 alone spheroids were then utilised in immune-killing assays, culturing with unstimulated or reovirus-stimulated PBMC. The results demonstrate that, in this more complex *in vitro* model, incorporating MSCs and TAMs, MG-63 cells exhibit an increased degree of resistance to immune-mediated killing by both unstimulated and OV- stimulated PBMC, compared with MG-63 cells alone in monolayer or spheroid culture (Figure 4B).

**Figure 4:**
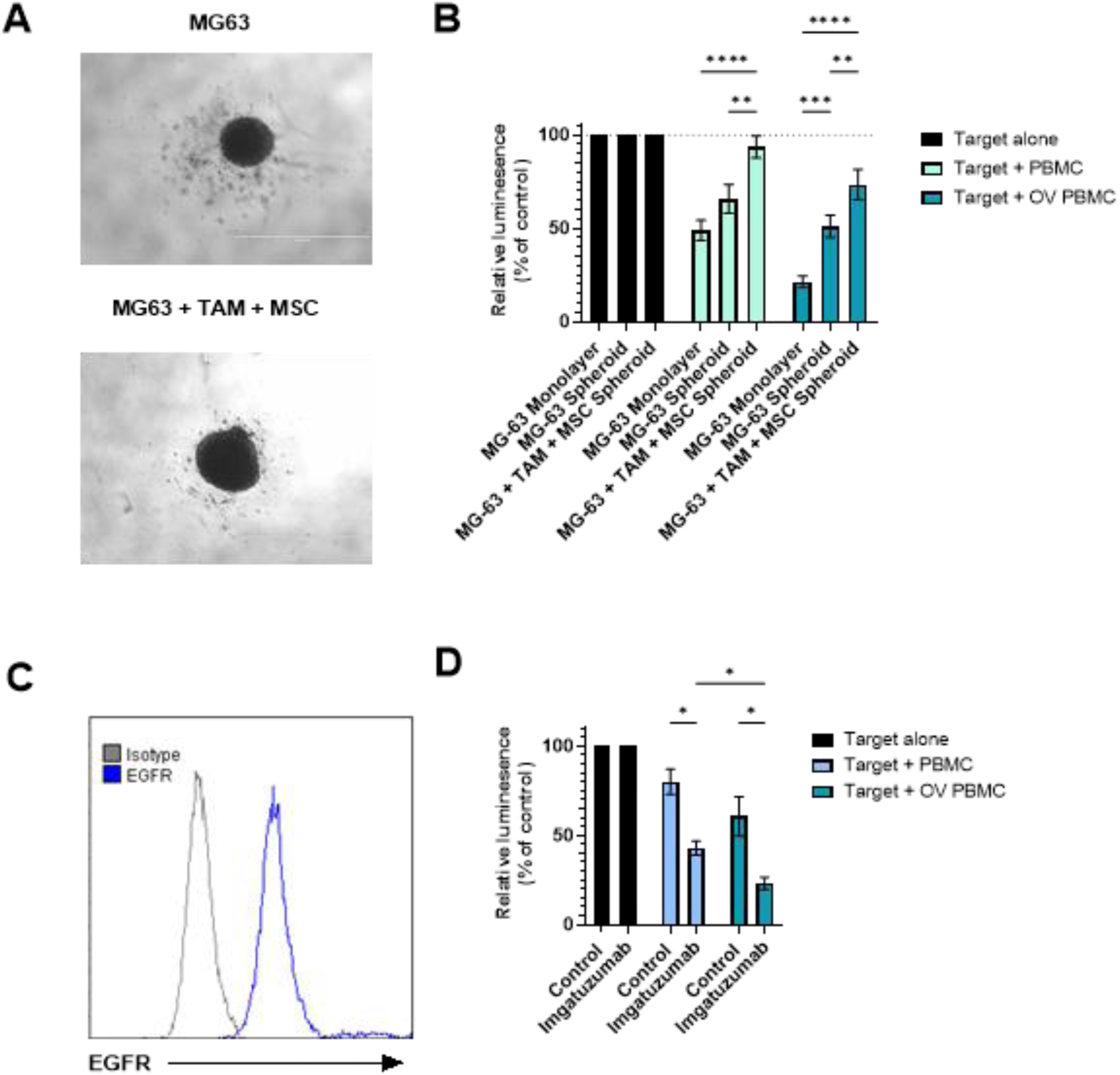
Immune-resistant multicellular spheroid model of OS. (A) OS spheroids were generated by seeding MG-63-LUC2 cell line alone or MG-63-Luc2 cells with CD14+ monocytes and MSCs at a ratio of 10:5:1 into low adhesion plates for 7 days. Spheroids were imaged using EVOS microscope at 4x magnification, scale bar = 1000 µm. (B) PBMC from healthy donors were treated ± reovirus at 0.1 PFU/cell for 48 hours, and then co-cultured at a ratio of 20:1 with MG-63-Luc2 cells in monolayer culture, spheroid culture or in multicellular spheroid co-culture (MG-63 + TAM + MSC) for 24 hours. D-luciferin was added to wells at 150 µg/mL and luminescence measured using a plate reader. Results presented as luminescence relative to none co-cultured target cell controls. Statistical comparison performed using two-way ANOVA, n=6. (C) EGFR expression on MG-63 cells was examined by staining with fluorescently conjugated antibody or matched isotype control and flow cytometry, representative histogram for n=3 experiments. (D) PBMC from healthy donors were treated ± reovirus at 0.1 PFU/cell for 48 hours/. Spheroid co-cultures (MG-63-Luc2 + TAM + MSC) were pre-treated with Imgatuzumab (anti-EGFR) at 10 µg/mL for 30 minutes. PBMC were co-cultured with spheroids at a ratio of 20:1 for 24 hours. D-luciferin was added to wells at 150 µg/mL and luminescence measured using a plate reader. Results presented as luminescence relative to none co-cultured target cell controls. Statistically analysed using two-way ANOVA, n=3.

### 3.5 Combining OV and antibody dependent cellular cytotoxicity (ADCC) to target OS

Monoclonal antibody (mAb)-based therapies are widely used and very successful in many cancers. An anti-EGFR antibody (cetuximab), underwent clinical testing in sarcoma patients but yielded disappointing results as a monotherapy (Ha et al., 2013). Since then, anti-EGFR monoclonal antibodies have been further developed, including the generation of a glycosylated mAb, Imgatuzumab for improved ADCC (Gerdes et al., 2013).

The increased resistance to immune mediated killing observed with the multicellular spheroid model provided an opportunity to test the combination of OV-activated PBMC with an anti-EGFR antibody (Imgatuzumab) using a more representative human OS tumour model. Critically, the MG-63 cell line used in the more complex, immunosuppressive multicellular spheroid model expressed EGFR (Figure 4C). PBMC were activated ± reovirus for 48 hours, and spheroids containing MG-63, TAMs and MSCs were pretreated with Imgatuzumab for 30 minutes. PBMCs were then added to spheroids at ratios of 20:1 and cultured for 24 hours. Importantly, combination of OV-stimulation of PBMC with Imgatuzumab significantly increased immune-mediated killing of MG-63 cells in immune-resistant spheroid cultures, when compared with either treatment alone (Figure 4D).

These data provide a proof of principle that ADCC of immune resistant OS tumours can be enhanced by the addition of OV. However, EGFR is expressed at varying levels and is absent in some OS tumours, therefore additional target antigens would need to be identified to offer alternative, personalised combination approaches (Lee et al., 2012).

### 3.6 Personalised mAb therapy in combination with HSV1716-GMCSF enhances immune-mediated killing of OS primary cells in immune-resistant multicellular spheroid models

To identify additional antigens targetable with mAbs, we analysed publicly available TARGET-OS bulk mRNA sequencing data from 88 OS patients. This confirmed that EGFR expression is heterogenous across OS patients. However, other potential candidate target antigens were identified, which included HER2 (ERBB2) and GD2 (B4GALNT1) (Figure 5A). Importantly, these antigens have an associated therapeutic mAb already used in the treatment of breast cancer and neuroblastoma respectively (Jeyakumar and Younis, 2012; Mora et al., 2025). First, we examined the expression of candidate target antigens in primary patient derived cell cultures. EGFR, GD2 and HER2 were all expressed, albeit to varying levels on both SARC011 and SARC050 cells (Figure 5B).

**Figure 5:**
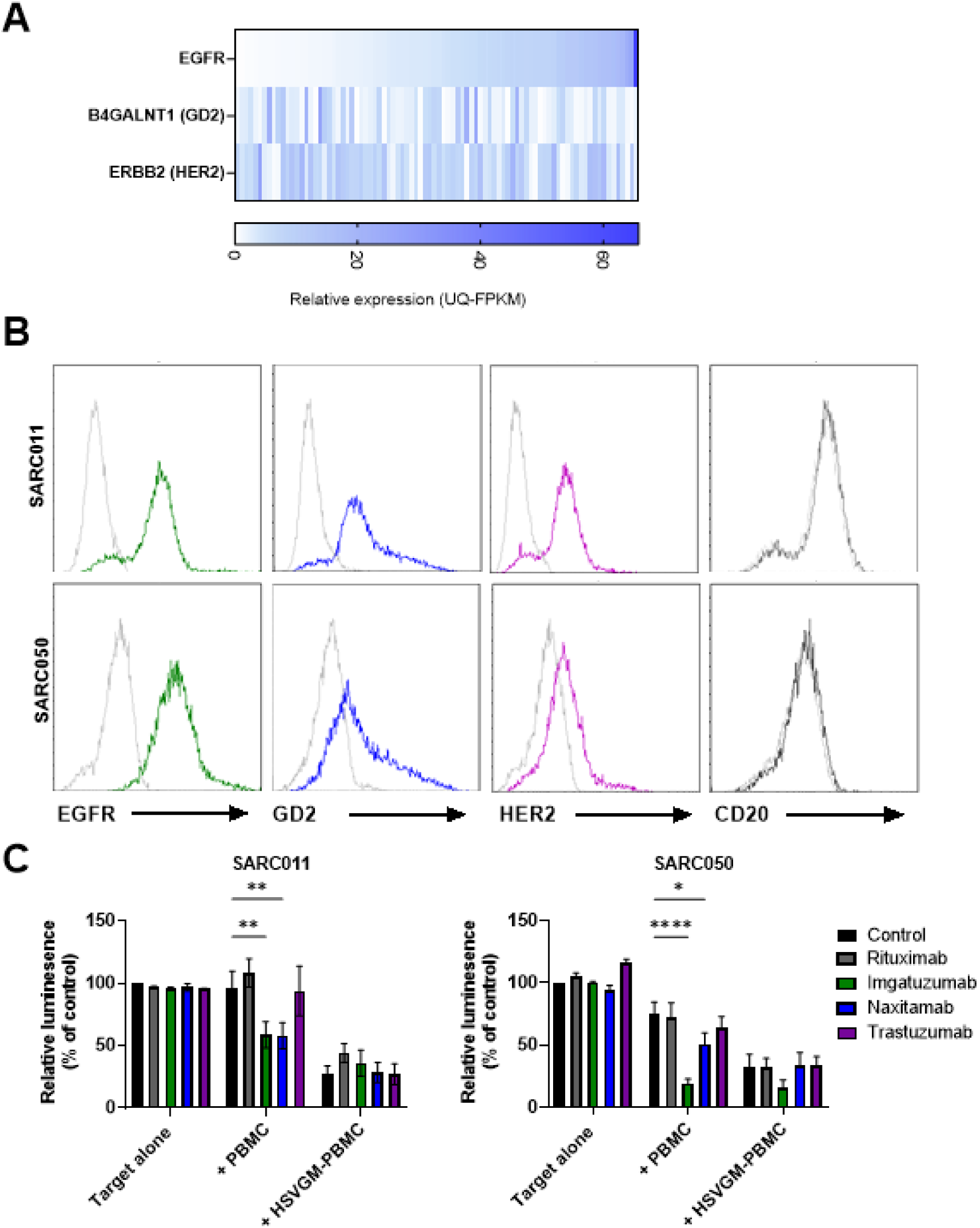
Monoclonal antibody treatment induces immune-mediated killing of OS primary cell cultures. (A) TARGET-OS bulk mRNA sequencing gene expression data (UQ-FPKM) from 88 OS patient tumours was downloaded for EGFR, B4GALNT1 (encoding a GD2 synthase enzyme) and ERBB2 (HER2) from NCI Genomic Data Commons (GDC; dbGaP phs000218). The data is ranked by expression of EGFR in the 88 tumour samples (lowest on left to highest on right). (B) SARC011 and SARC050 cells were stained with fluorescently conjugated anti-EGFR, anti-GD2, anti-HER2 or anti-CD20 antibodies or matched isotype controls, expression was assessed by flow cytometry, histogram overlays showing surface protein expression (colour) and isotype control (grey). Representative of n=3 independent experiments. (C) Healthy donor PBMCs were treated ± HSV1716 GMCSF at 0.1 PFU/cell for 48 hours. SARC011-Luc2 or SARC050-Luc2 cells were cultured as monolayers and treated with Rituximab (anti-CD20), Imgatuzumab (anti-EGFR), Naxitamab (anti-GD2) or Trastuzumab (anti-HER2) at 10 µg/ mL for 30 minutes. Unstimulated or HSV1716-GMCSF activated PBMCs were then added to cells at a ratio of 20:1 for 24 hours. D-luciferin was added to wells at 150 µg/mL and luminescence measured using a plate reader. Results presented as mean luminescence relative to untreated target cells ± SEM for n=3 independent experiments. Statistical comparison performed using two-way ANOVA.

Next, we performed immune killing assays using SARC011 and SARC050 target cells in monolayer cultures, to compare the efficacy of each mAb. To do this, we combined HSV1716-GMCSF activation of PBMC with mAb pre-treatment of OS cells, targeting either EGFR (Imgatuzumab), GD2 (Naxitamab) or HER2 (Trastuzumab). Rituximab, a chimaeric antibody targeting CD20, which is not expressed on OS primary cell cultures (Figure 5B), served as a negative control to confirm that any enhancement of immune-mediated killing was dependent on tumour antigen expression. In both SARC011 and SARC050 cells, Imgatuzumab and Naxitamab treatment enhanced immune-mediated killing by unstimulated PBMCs (Figure 5C). However, combination of HSV1716-GMCSF activation of PBMC and mAb treatment had no apparent additive/ synergistic effect, likely due to the fact that HSV1716-GMCSF activation of PBMC alone is highly effective in these more simplistic monolayer models. Therefore, we proceeded to test the efficacy of combined HSV1716- GMCSF activation of PBMC and mAb treatment in SARC011 and SARC050 spheroids, either with primary cell cultures alone or cultured with MSCs and CD14+ monocytes.

Immune killing assays were carried out to test combinations of HSV1716-GMCSF based activation of PBMC and mAb pretreatment of target cells. In SARC011 spheroids, both Imgatuzumab and Naxitamab significantly increased immune-mediated killing by HSV1716- GMCSF stimulated PBMCs (Figure 6A). However, for SARC050 cells, mAb treatment only enhanced immune-mediated killing by unstimulated PBMC (Figure 6A). Importantly, the more complex multicellular immune-resistant spheroid model, incorporating primary OS cells, MSC and TAM, was markedly more resistant to HSV1716-GMCSF activated immune-mediated killing when compared with monolayer and spheroid cultures (Figure 5C and 6). However, combining HSV1716-GMCSF activation of PBMCs with either Naxitamab for SARC011 cells and Imgatuzumab for SARC050 cells significantly increased immune-mediated killing (Figure 6B), indicating that more complex *in vitro* model systems are required to screen effective combination strategies to improve immune-mediated killing of OS.

**Figure 6:**
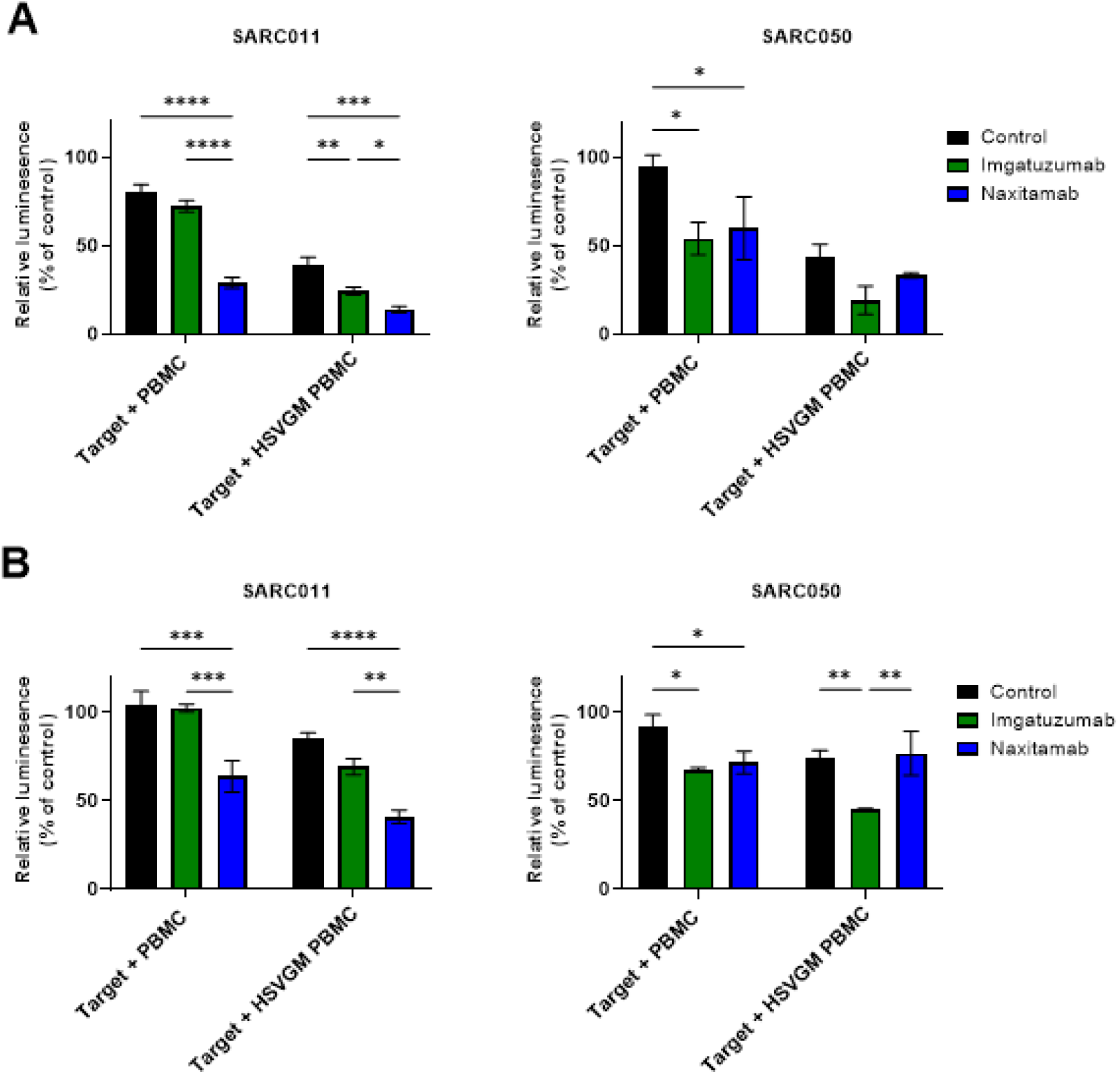
Combination of HSV1716-GMCSF and targeted mAb enhances immune-mediated destruction of immune-resistant OS multicellular spheroids. (A-B) Healthy donor PBMCs were treated ± HSV1716 GMCSF at 0.1 PFU/cell for 48 hours. SARC011-Luc2 or SARC050-Luc2 cells were cultured as spheroids in (A) monoculture or (B) multicellular cultures containing TAMs and MSCs. Cells were treated with a control antibody (Rituximab; anti-CD20), Imgatuzumab (anti-EGFR) or Naxitamab (anti-GD2) at 10 µg/ mL for 30 minutes. Unstimulated or HSV1716-GMCSF activated PBMCs were then added to cells at a ratio of 20:1 for 24 hours. D-luciferin was added to wells at 150 µg/mL and luminescence measured using a plate reader. Results presented as mean luminescence relative to untreated target cells ± SEM for n=3 independent experiments. Statistical comparison performed using two-way ANOVA.

## 4. Discussion

We demonstrate that multiple oHSVs, including HSV1716, HSV1716-GMCSF and HSV47Δ exert potent direct oncolytic activity against OS cell lines, primary patient-derived cultures, and OS cells from freshly resected tumour samples. Moreover, HSV1716-GMCSF treatment was demonstrated to repolarise *in vitro* generated OS TAMs to a similar extent as Mifamurtide, a clinically approved OS treatment. Finally, in a multicellular spheroid model of OS (incorporating primary OS cell cultures, MSCs and TAMS), HSV1716-GMCSF in combination with targeted therapeutic antibodies improved immune-mediated destruction of OS. In summary, these findings indicate that HSV1716-GMCSF in combination with targeted mAbs antibodies may offer a valuable approach to OS immunotherapy, where HSV1716- GMCSF induces direct oncolysis, repolarises TAMs and activates immune-mediated killing of OS, and combination with mAbs enhances immune mediated killing by ADCC.

Previous studies have reported the oncolytic effects of several oHSVs (including HSVN212- eGFP, G207 and NV1020) against long-established OS cell lines (Bharatan et al., 2002; Boeuf et al., 2017). Here we report a comparison of three distinct oHSVs across several human OS cell types, including primary OS cells more recently established from patient tumours. Our study presents the most comprehensive analysis of OV activity in human OS preclinical model systems to date. Each of the oHSVs exhibited significant levels of direct oncolytic activity against all OS monolayer cultures after 7 days of treatment. However, HSV47Δ had superior oncolytic activity when compared with HSV1716 and HSV1716- GMCSF in some cell cultures, and this was consistent with significantly increased replication of HSV47Δ in HOS, MG-63 and SARC050 cells when compared with other oHSVs. The deletion of the α*47* gene in HSV47Δ is known to enhance oHSV replication in tumour cells (Todo et al., 2001). Importantly, α*47* is upstream of *US11* in the viral genome, and its deletion causes promoter de-repression, leading to early expression of *US11*, a potent protein kinase R (PKR) antagonist, allowing the virus to rapidly neutralise PKR mediated antiviral signalling and providing HSV47Δ with a replication advantage in tumour cells with partially intact PKR pathways (Todo et al., 2001). In addition, we also observed that HSV47Δ treatment of primary cell cultures resulted in syncytia formation. Several human herpes simplex viruses have been reported to form syncytia, and the fusion of cell membranes, improves viral spread and replication, which may also account for increased early replication of this virus (Burton and Bartee, 2019; Yan et al., 2025).

Importantly, the oncolytic activity of oHSVs represents only one of their multimodal mechanisms of action. The OV field has shifted significantly in recent years from viewing OVs as direct oncolytic agents to recognising that immune activation and immunogenic cell death are central to their therapeutic efficacy (Inoue, 2026). Among the oHSVs tested, HSV1716-GMCSF demonstrated the strongest immune-stimulatory profile; HSV1716- GMCSF treatment of PBMC induced a type-I IFN response and production of cytokines which activate and polarise macrophage. Although not tested here, we have previously shown that OV induced IFN-I responses activate human NK cells *in vitro* and that the peak of IFN-I responses and NK cell activation are co-incidental in patients *in vivo* (Parrish et al., 2015; El-Sherbiny et al., 2015; Wantoch et al., 2022). It seems likely that the HSV1716- GMCSF analysed here activates NK cells via IFN-I responses and modulates macrophage polarisation via the encoded GM-CSF and induction of other cytokines such as IL-6 and IL- 1β. TAMs are recognised as central drivers of immunosuppression and progression in OS, and strategies to reprogramme them towards a pro-inflammatory, anti-tumour phenotype are a major focus of current research (Luo et al., 2020). The ability of HSV1716-GMCSF to modulate TAMs aligns with emerging data that GM-CSF-armed OVs can enhance myeloid activation and antigen presentation, thereby amplifying downstream adaptive and innate immune responses (Jennings et al., 2019; Tijtgat et al., 2021; Ageenko et al., 2026). Moreover, oHSVs are increasingly used as cancer selective gene delivery vectors, enabling intratumoral expression of cytokines, chemokines, or other regulatory molecules such as microRNAs to enhance immune activation and/ or remodel the immunosuppressive TME (Jennings et al., 2024; Wang et al., 2024).

This work demonstrates that HSV1716-GMCSF treatment may be superior to or least complement the action of Mifamurtide. Mifamurtide is delivered in liposomes and phagocytosed by myeloid-lineage cells after intravenous administration. The liposomal vesicles are degraded and the active component (muramyl tripeptide phosphatidylethanolamine; MTP-PE) binds nucleotide-binding oligomerization domain (NOD)-2 receptors, activating myeloid cells, and stimulating secretion of inflammatory cytokines including IL-1β and IL-6, leading to immune activation (Fogler and Fidler, 1987;

Ando et al., 2011). However, Mifamurtide has no direct stimulatory action on cytotoxic immune cells such as NK cells. While both HSV1716-GMCSF and Mifamurtide were capable of TAM repolarisation, the unique ability of HSV1716-GMCSF to simultaneously activate NK cells and stimulate immune-mediated killing of OS demonstrates potential benefits of the OV-based therapy.

A phase I/II clinical trial recently assessed the efficacy of intratumoral OH2 (an HSV type 2 strain HG52 expressing GM-CSF) in locally advanced or metastatic sarcoma. This trial included four bone sarcoma patients (OS or Ewing sarcoma). The study evaluated OH2 alone or in combination with PD-1 blockade (antibody HX008), and demonstrated that the monotherapy and combination were safe and well tolerated in patients, warranting further investigation (Tan et al., 2025). Importantly, this study demonstrates the safety of oHSV encoding GM-CSF in sarcoma patients. The clinically approved OV, Talimogene Laherparepvec (T-Vec) also expresses GM-CSF and is used in the treatment of melanoma (O’Donoghue et al., 2016).

Ringwalt *et al* recently reported that combining oHSV with the alkylating agent trabectedin reduced tumour burden in syngeneic mice harbouring K7M2 and F420 murine OS tumours (Ringwalt et al., 2024). Critically, depletion of NK cells from these OS bearing mice diminished the synergistic effects of oHSV with trabectedin. Interestingly, as well as having direct anti-tumour activity, trabectedin modulates the TME, in particular the myeloid cell compartment (D’Incalci et al, 2014), suggesting that activation of NK cells in this treatment combination might occur via the OV (e.g. by the action of IFN-I) and by trabectedin-mediated reduction in immunosuppression. Although murine sarcoma models permit evaluation of oHSVs in immunocompetent hosts, differences between mouse and human immune systems and mouse and human sarcoma biology make validation in representative human models essential.

To better capture the complexity of the OS TME in human *in vitro* models, we employed a multicellular spheroid model incorporating OS cells, MSCs and TAMs, which exhibited resistance to immune-mediated killing, more closely reflecting the immunosuppressive milieu observed in patients. The use of such 3D, multicellular systems is increasingly recognised as critical for preclinical evaluation of immunotherapies in solid tumours, as they more accurately recapitulate cell–cell interactions and immunosuppressive TMEs than traditional 2D cultures (Zhou et al., 2024). Pairing HSV1716-GMCSF with anti-GD2 or anti-EGFR mAbs (selected according to tumour antigen expression) significantly increased immune-mediated killing in our multicellular spheroid models. Therapeutic antibodies based on IgG an engage NK cells via the low affinity Fc receptor CD16 and induce ADCC; our data suggest that oHSV-mediated NK cell activation synergises with therapeutic antibody targeting to enhance tumour cell killing. The anti-EGFR antibody cetuximab has previously been described to induce NK cell–mediated ADCC against EGFR-expressing osteosarcoma cell lines (Pahl et al., 2012). However, the monolayer cell line cultures used in this study fail to represent the complexities of the OS TME (such as the immunosuppressive environment), and the potential need for combination strategies. The efficacy of combining anti-GD2 mAb with recombinant GM-CSF treatment has been demonstrated in neuroblastoma (Yu et al., 2010; Voeller and Sodel, 2019). Critically, a phase III clinical trial in high-risk neuroblastoma patients comparing standard isotretinoin therapy or combination with GMCSF, anti-GD2 mAb and IL-2 increased 2-year overall survival from 75% to 86% (Yu et al., 2010).

Importantly, oHSVs have demonstrated safety and efficacy profiles in other cancers, resulting in their approval for melanoma and glioblastoma (Zheng et al., 2025). Moreover, their safety profile in paediatric cancer patients (including a total of 4 OS patients) has been demonstrated using both intravenous and intratumoral administration in phase I clinical trials (Streby et al., 2017; Streby et al., 2019). From a translational perspective, the use of HSV1716-GMCSF in combination with clinically available therapeutic antibodies is particularly attractive, withoHSVs, anti-GD2 and anti-EGFRantibodies already approved for use in other cancers. When paired with therapeutic antibodies, HSV1716-GMCSF offers a personalised strategy which integrates direct oncolysis, enhanced immune-mediated tumour destruction, and remodelling of the immunosuppressive TME. Given the limited clinical progress in OS immunotherapy to date, these data provide a strong rationale for further investigation of oHSV–mAb combination strategies.

## Acknowledgments

The authors thank the Bone Cancer Research Trust for their generous funding and support of this work. We acknowledge support from the CRUK Experimental Cancer Centre, the CRUK City of London Centre Award [CTRQQR-2021\100004]. We thank the UCL/UCLH Biobank for Studying Health and Disease and the Research Innovation Centre at RNOH and ROH Birmingham for provision of human tissue samples and clinical data. We thank the patients for the generous donation of their material without which this research would not have been possible. We also extend our thanks to the additional organisations whose contributions supported this study.

## Author contributions

**TB:** Data curation, Formal analysis, Funding acquisition, Investigation, Methodology, Project administration, Resources, Supervision, Validation, Visualisation, Writing – original draft, review & editing. **LC:** Methodology, Resources, Writing – review & editing. **KS:** Investigation, Writing – review & editing. **VAJ:** Investigation, Methodology, Resources, Supervision, Writing – review & editing. **SD:** Investigation, Data curation, Formal analysis, Writing – review & editing. **RTB:** Investigation, Writing – review & editing. **JS:** Investigation, Writing – review & editing. **HP:** Resources, Writing – review & editing. **GS:** Resources, Writing – review & editing. **JuS:** Methodology, Resources, Writing - review and editing**. HO:** Resources, Writing – review & editing. **DM:** Resources, Writing – review & editing. **PVG:** Resources, Writing – review & editing. **NJC:** Investigation, Methodology, Resources, Supervision, Writing – review & editing. **SS:** Resources, Methodology, Writing – review & editing. **FEM:** Conceptualisation, Funding acquisition, Supervision, Investigation, Methodology, Project administration, Writing – original draft, review & editing. **GPC:** Conceptualisation, Funding acquisition, Supervision, Investigation, Methodology, Project administration, Writing – original draft, review & editing.

## References

Ageenko, A.B., Vasileva, N.S., Chesnokova, A.S., Semenov, D. V., Byvakina, A.A., Dymova, M.A., Sen’kova, A. V., Nushtaeva, A.A., Leonteva, A.A., Savinovskaya, Y.I., Kochneva, G. V., Richter, V.A. and Kuligina, E. V. 2026. Oncolytic Virus VV-GMCSF-Lact and Human GM-CSF Against GL261 Glioma in Immunocompetent Mice. Pharmaceuticals. 19(3), p.434.

Anderson, P.M., Meyers, P., Kleinerman, E., Venkatakrishnan, K., Hughes, D.P., Herzog, C., Huh, W., Sutphin, R., Vyas, Y.M., Shen, V., Warwick, A., Yeager, N., Oliva, C., Wang, B., Liu, Y. and Chou, A. 2014. Mifamurtide in metastatic and recurrent osteosarcoma: a patient access study with pharmacokinetic, pharmacodynamic, and safety assessments. Pediatric blood & cancer. 61(2), pp.238–44.

Ando, K., Mori, K., Corradini, N., Redini, F. and Heymann, D. 2011. Mifamurtide for the treatment of nonmetastatic osteosarcoma. Expert opinion on pharmacotherapy. 12(2), pp.285–92.

Andtbacka, R.H.I., Ross, M., Puzanov, I., Milhem, M., Collichio, F., Delman, K.A., Amatruda, T., Zager, J.S., Cranmer, L., Hsueh, E., Chen, L., Shilkrut, M. and Kaufman, H.L. 2016. Patterns of Clinical Response with Talimogene Laherparepvec (T-VEC) in Patients with Melanoma Treated in the OPTiM Phase III Clinical Trial. Annals of Surgical Oncology. 23(13), pp.4169–4177.

Andreatta, M., & Carmona, S. J. (2021). UCell: Robust and scalable single-cell gene signature scoring. Computational and structural biotechnology journal, 19, 3796–3798.

Barr, T., Jennings, V.A., Roundhill, E.A., Baugh, R.T., Yamrali, M., Owston, H.E., McGonagle, D., Giannoudis, P. V., Caplen, N.J., Khan, J., Bell, J.C., Burchill, S.A., Errington-Mais, F. and Cook, G.P. 2025. Oncolytic Maraba Virus MG1 Mediates Direct and Natural Killer Cell-Dependent Lysis of Ewing Sarcoma. Cancers. 17(20), p.3319.

Bharatan, N.S., Currier, M.A. and Cripe, T.P. 2002. Differential susceptibility of pediatric sarcoma cells to oncolysis by conditionally replication-competent herpes simplex viruses. Journal of Pediatric Hematology/Oncology. 24(6), pp.447–453.

Boeuf, F. Le, Selman, M., Son, H.H., Bergeron, A., Chen, A., Tsang, J., Butterwick, D., Arulanandam, R., Forbes, N.E., Tzelepis, F., Bell, J.C., Werier, J., Abdelbary, H. and Diallo, J. 2017. Oncolytic Maraba Virus MG1 as a Treatment for Sarcoma. International Journal of Cancer. 141(6), pp.1257–1264.

Bouzeineddine, N.Z., Philippi, A., Gee, K. and Basta, S. 2025. Granulocyte macrophage colony stimulating factor in virus-host interactions and its implication for immunotherapy. Cytokine & Growth Factor Reviews. 81, pp.54–63.

Burton, C. and Bartee, E. 2019. Syncytia Formation in Oncolytic Virotherapy. Molecular therapy oncolytics. 15, pp.131–139.

Cao, F., Nguyen, P., Hong, B., DeRenzo, C., Rainusso, N.C., Rodriguez Cruz, T., Wu, M.-F., Liu, H., Song, X.-T., Suzuki, M., Wang, L.L., Yustein, J.T. and Gottschalk, S. 2021. Engineering Oncolytic Vaccinia Virus to redirect Macrophages to Tumor Cells. Advances in cell and gene therapy. **4**(2).

Currier, M.A., Eshun, F.K., Sholl, A., Chernoguz, A., Crawford, K., Divanovic, S., Boon, L., Goins, W.F., Frischer, J.S., Collins, M.H., Leddon, J.L., Baird, W.H., Haseley, A., Streby, K.A., Wang, P.-Y., Hendrickson, B.W., Brekken, R.A., Kaur, B., Hildeman, D. and Cripe, T.P. 2013. VEGF Blockade Enables Oncolytic Cancer Virotherapy in Part by Modulating Intratumoral Myeloid Cells. Molecular Therapy. 21(5), pp.1014–1023.

D’Incalci, M., Badri, N., Galmarini, C. M., & Allavena, P. (2014). Trabectedin, a drug acting on both cancer cells and the tumour microenvironment. British journal of cancer, 111(4), 646–650.

Eghtedari, A.R., Vaezi, M.A., Safari, E., Salimi, V., Safizadeh, B., Babaheidarian, P., Abiri, A., Mahdinia, E., Alireza Mirzaei, Mokhles, P. and Tavakoli-Yaraki, M. 2023. The expression changes of PD-L1 and immune response mediators are related to the severity of primary bone tumors. Scientific Reports. 13(1), p.20474.

El-Sherbiny, Y. M., Holmes, T. D., Wetherill, L. F., Black, E. V., Wilson, E. B., Phillips, S. L., Scott, G. B., Adair, R. A., Dave, R., Scott, K. J., Morgan, R. S., Coffey, M., Toogood, G. J., Melcher, A. A., & Cook, G. P. (2015). Controlled infection with a therapeutic virus defines the activation kinetics of human natural killer cells in vivo. Clinical and Experimental Immunology, 180(1), 98–107.

Ferrari, S., Briccoli, A., Mercuri, M., Bertoni, F., Picci, P., Tienghi, A., Del Prever, A.B., Fagioli, F., Comandone, A. and Bacci, G. 2003. Postrelapse Survival in Osteosarcoma of the Extremities: Prognostic Factors for Long-Term Survival. Journal of Clinical Oncology. 21(4), pp.710–715.

Fogler, W.E. and Fidler, I.J. 1987. Comparative interaction of free and liposome-encapsulated nor-muramyl dipeptide or muramyl tripeptide phosphatidylethanolamine (3H-labelled) with human blood monocytes. International Journal of Immunopharmacology. 9(2), pp.141–150.

Gerdes, C.A., Nicolini, V.G., Herter, S., van Puijenbroek, E., Lang, S., Roemmele, M., Moessner, E., Freytag, O., Friess, T., Ries, C.H., Bossenmaier, B., Mueller, H.J. and Umaña, P. 2013. GA201 (RG7160): a novel, humanized, glycoengineered anti-EGFR antibody with enhanced ADCC and superior in vivo efficacy compared with cetuximab. Clinical Cancer Research 19(5), pp.1126–38.

Gerrand, C., Amary, F., Anwar, H.A., Brennan, B., Dileo, P., Kalkat, M.S., McCabe, M.G., McCullough, A.L., Parry, M.C., Patel, A., Seddon, B.M., Sherriff, J.M., Tirabosco, R. and Strauss, S.J. 2025. UK guidelines for the management of bone sarcomas. British Journal of Cancer. 132(1), pp.32–48.

Ha, H.T., Griffith, K.A., Zalupski, M.M., Schuetze, S.M., Thomas, D.G., Lucas, D.R., Baker, L.H. and Chugh, R. 2013. Phase II Trial of Cetuximab in Patients With Metastatic or Locally Advanced Soft Tissue or Bone Sarcoma. American Journal of Clinical Oncology. 36(1), pp.77–82.

Han, Z., Chen, G. and Wang, D. 2025. Emerging immunotherapies in osteosarcoma: from checkpoint blockade to cellular therapies. Frontiers in Immunology. 16:1579822.

Hawkins, A. G., Shapiro, J. A., Spielman, S. J., Mejia, D. S., Venkatesh Prasad, D., Ichihara, N., Yakovets, A., Gottlieb, A. M., Wheeler, K. G., Bethell, C. J., Foltz, S. M., O’Malley, J., Greene, C. S., & Taroni, J. N. (2026). The Single-Cell Pediatric Cancer Atlas: Data portal and open-source tools for single-cell transcriptomics of pediatric tumors. Cell genomics, 6(7), 101283.

Heath, A.P., Ferretti, V., Agrawal, S., An, M., Angelakos, J.C., Arya, R., Bajari, R., Baqar, B., Barnowski, J.H.B., Burt, J., Catton, A., Chan, B.F., Chu, F., Cullion, K., Davidsen, T., Do, P.-M., Dompierre, C., Ferguson, M.L., Fitzsimons, M.S., Ford, M., Fukuma, M., Gaheen, S., Ganji, G.L., Garcia, T.I., George, S.S., Gerhard, D.S., Gerthoffert, F., Gomez, F., Han, K., Hernandez, K.M., Issac, B., Jackson, R., Jensen, M.A., Joshi, S., Kadam, A., Khurana, A., Kim, K.M.J., Kraft, V.E., Li, S., Lichtenberg, T.M., Lodato, J., Lolla, L., Martinov, P., Mazzone, J.A., Miller, D.P., Miller, I., Miller, J.S., Miyauchi, K., Murphy, M.W., Nullet, T., Ogwara, R.O., Ortuño, F.M., Pedrosa, J., Pham, P.L., Popov, M.Y., Porter, J.J., Powell, R., Rademacher, K., Reid, C.P., Rich, S., Rogel, B., Sahni, H., Savage, J.H., Schmitt, K.A., Simmons, T.J., Sislow, J., Spring, J., Stein, L., Sullivan, S., Tang, Y., Thiagarajan, M., Troyer, H.D., Wang, C., Wang, Z., West, B.L., Wilmer, A., Wilson, S., Wu, K., Wysocki, W.P., Xiang, L., Yamada, J.T., Yang, L., Yu, C., Yung, C.K., Zenklusen, J.C., Zhang, J., Zhang, Z., Zhao, Y., Zubair, A., Staudt, L.M. and Grossman, R.L. 2021. The NCI Genomic Data Commons. Nature Genetics. 53(3), pp.257–262.

Hu, C., Li, T., Xu, Y., Zhang, X., Li, F., Bai, J., Chen, J., Jiang, W., Yang, K., Ou, Q., Li, X., Wang, P., & Zhang, Y. (2023). CellMarker 2.0: an updated database of manually curated cell markers in human/mouse and web tools based on scRNA-seq data. Nucleic acids research, 51(D1), D870–D876.

Inoue, H. 2026. Oncolytic Virotherapy and Immunogenic Cell Death: Mechanisms, Platforms, and Clinical Translation. Viruses. 18(4), p.461.

Jennings, V.A., Rumbold-Hall, R., Migneco, G., Barr, T., Reilly, K., Ingram, N., St Hilare, I., Heaton, S., Alzamel, N., Jackson, D., Ralph, C., Banerjee, S., McNeish, I., Bell, J.C., Melcher, A.A., Ilkow, C., Cook, G.P. and Errington-Mais, F. 2024. Enhancing oncolytic virotherapy by extracellular vesicle mediated microRNA reprograming of the tumour microenvironment. Frontiers in immunology. 15, p.1500570.

Jennings, V.A., Scott, G.B., Rose, A.M.S., Scott, K.J., Migneco, G., Keller, B., Reilly, K., Donnelly, O., Peach, H., Dewar, D., Harrington, K.J., Pandha, H., Samson, A., Vile, R.G., Melcher, A.A. and Errington-Mais, F. 2019. Potentiating Oncolytic Virus-Induced Immune-Mediated Tumor Cell Killing Using Histone Deacetylase Inhibition. Molecular Therapy. 27(6), pp.1139–1152.

Jeyakumar, A. and Younis, T. 2012. Trastuzumab for HER2-Positive Metastatic Breast Cancer: Clinical and Economic Considerations. Clinical Medicine Insights. Oncology. 6, pp.179–87.

Kager, L., Zoubek, A., Pötschger, U., Kastner, U., Flege, S., Kempf-Bielack, B., Branscheid, D., Kotz, R., Salzer-Kuntschik, M., Winkelmann, W., Jundt, G., Kabisch, H., Reichardt, P., Jürgens, H., Gadner, H. and Bielack, S.S. 2003. Primary Metastatic Osteosarcoma: Presentation and Outcome of Patients Treated on Neoadjuvant Cooperative Osteosarcoma Study Group Protocols. Journal of Clinical Oncology. 21(10), pp.2011– 2018.

Kaur, B., Antonio Chiocca, E. and P. Cripe, T. 2012. Oncolytic HSV-1 Virotherapy: Clinical Experience and Opportunities for Progress. Current Pharmaceutical Biotechnology. 13(9), pp.1842–1851.

Lee, J.A., Ko, Y., Kim, D.H., Lim, J.S., Kong, C.-B., Cho, W.H., Jeon, D.-G., Lee, S.-Y. and Koh, J.-S. 2012. Epidermal growth factor receptor: is it a feasible target for the treatment of osteosarcoma? Cancer research and treatment. 44(3), pp.202–9.

Luo, Z.-W., Liu, P.-P., Wang, Z.-X., Chen, C.-Y. and Xie, H. 2020. Macrophages in Osteosarcoma Immune Microenvironment: Implications for Immunotherapy. Frontiers in oncology. 10, p.586580.

Lussier, D.M., Johnson, J.L., Hingorani, P. and Blattman, J.N. 2015. Combination immunotherapy with α-CTLA-4 and α-PD-L1 antibody blockade prevents immune escape and leads to complete control of metastatic osteosarcoma. Journal for ImmunoTherapy of Cancer. 3(1), p.21.

Meyers, P.A., Schwartz, C.L., Krailo, M.D., Healey, J.H., Bernstein, M.L., Betcher, D., Ferguson, W.S., Gebhardt, M.C., Goorin, A.M., Harris, M., Kleinerman, E., Link, M.P., Nadel, H., Nieder, M., Siegal, G.P., Weiner, M.A., Wells, R.J., Womer, R.B., Grier, H.E. and Children’s Oncology Group 2008. Osteosarcoma: the addition of muramyl tripeptide to chemotherapy improves overall survival--a report from the Children’s Oncology Group. Journal of clinical oncology : official journal of the American Society of Clinical Oncology. 26(4), pp.633–8.

Mirabello, L., Troisi, R.J. and Savage, S.A. 2009. International osteosarcoma incidence patterns in children and adolescents, middle ages and elderly persons. International journal of cancer. 125(1), pp.229–34.

Mora, J., Chan, G.C.F., Morgenstern, D.A., Amoroso, L., Nysom, K., Faber, J., Wingerter, A., Bear, M.K., Rubio-San-Simon, A., de Las Heras, B.M., Tornøe, K., Düring, M. and Kushner, B.H. 2025. The anti-GD2 monoclonal antibody naxitamab plus GM-CSF for relapsed or refractory high-risk neuroblastoma: a phase 2 clinical trial. Nature Communications. 16(1), p.1636.

Munoz-Garcia, J., Jubelin, C., Loussouarn, A., Goumard, M., Griscom, L., Renodon-Cornière, A., Heymann, M.-F. and Heymann, D. 2021. In vitro three-dimensional cell cultures for bone sarcomas. Journal of Bone Oncology. 30, p.100379.

O’Donoghue, C., Doepker, M.P. and Zager, J.S. 2016. Talimogene laherparepvec: overview, combination therapy and current practices. Melanoma Management. 3(4), pp.267–272.

Pahl, J.H.W., Ruslan, S.E.N., Buddingh, E.P., Santos, S.J., Szuhai, K., Serra, M., Gelderblom, H., Hogendoorn, P.C.W., Egeler, R.M., Schilham, M.W. and Lankester, A.C. 2012. Anti-EGFR antibody cetuximab enhances the cytolytic activity of natural killer cells toward osteosarcoma. Clinical Cancer Research. 18(2), pp.432–441.

Parrish, C., Scott, G. B., Migneco, G., Scott, K. J., Steele, L. P., Ilett, E., West, E. J., Hall, K., Selby, P. J., Buchanan, D., Varghese, A., Cragg, M. S., Coffey, M., Hillmen, P., Melcher, A. A., & Errington-Mais, F. (2015). Oncolytic reovirus enhances rituximab-mediated antibody-dependent cellular cytotoxicity against chronic lymphocytic leukaemia. Leukemia, 29(9), 1799–1810.

Ringwalt, E.M., Currier, M.A., Glaspell, A.M., Chen, C.-Y., Cannon, M. V., Cam, M., Gross, A.C., Gust, M., Wang, P.-Y., Boon, L., Biederman, L.E., Schwarz, E., Rajappa, P., Lee, D.A., Mardis, E.R., Carson, W.E., Roberts, R.D. and Cripe, T.P. 2024. Trabectedin promotes oncolytic virus antitumor efficacy, viral gene expression, and immune effector function in models of bone sarcoma. Molecular Therapy: Oncology. 32(4), p.200886.

Stiller, C.A., Trama, A., Serraino, D., Rossi, S., Navarro, C., Chirlaque, M.D. and Casali, P.G. 2013. Descriptive epidemiology of sarcomas in Europe: Report from the RARECARE project. European Journal of Cancer. 49(3), pp.684–695.

Strauss, S.J., Frezza, A.M., Abecassis, N., Bajpai, J., Bauer, S., Biagini, R., Bielack, S., Blay, J.Y., Bolle, S., Bonvalot, S., Boukovinas, I., Bovee, J.V.M.G., Boye, K., Brennan, B., Brodowicz, T., Buonadonna, A., de Álava, E., Dei Tos, A.P., Garcia del Muro, X., Dufresne, A., Eriksson, M., Fagioli, F., Fedenko, A., Ferraresi, V., Ferrari, A., Gaspar, N., Gasperoni, S., Gelderblom, H., Gouin, F., Grignani, G., Gronchi, A., Haas, R., Hassan, A.B., Hecker-Nolting, S., Hindi, N., Hohenberger, P., Joensuu, H., Jones, R.L., Jungels, C., Jutte, P., Kager, L., Kasper, B., Kawai, A., Kopeckova, K., Krákorová, D.A., Le Cesne, A., Le Grange, F., Legius, E., Leithner, A., López Pousa, A., Martin-Broto, J., Merimsky, O., Messiou, C., Miah, A.B., Mir, O., Montemurro, M., Morland, B., Morosi, C., Palmerini, E., Pantaleo, M.A., Piana, R., Piperno-Neumann, S., Reichardt, P., Rutkowski, P., Safwat, A.A., Sangalli, C., Sbaraglia, M., Scheipl, S., Schöffski, P., Sleijfer, S., Strauss, D., Sundby Hall, K., Trama, A., Unk, M., van de Sande, M.A.J., van der Graaf, W.T.A., van Houdt, W.J., Frebourg, T., Ladenstein, R., Casali, P.G. and Stacchiotti, S. 2021. Bone sarcomas: ESMO–EURACAN–GENTURIS–ERN PaedCan Clinical Practice Guideline for diagnosis, treatment and follow-up. Annals of Oncology. 32(12), pp.1520–1536.

Streby, K.A., Currier, M.A., Triplet, M., Ott, K., Dishman, D.J., Vaughan, M.R., Ranalli, M.A., Setty, B., Skeens, M.A., Whiteside, S., Yeager, N.D., Haworth, K.B., Simpson, K., Conner, J. and Cripe, T.P. 2019. First-in-Human Intravenous Seprehvir in Young Cancer Patients: A Phase 1 Clinical Trial. Molecular therapy : the journal of the American Society of Gene Therapy. 27(11), pp.1930–1938.

Streby, K.A., Geller, J.I., Currier, M.A., Warren, P.S., Racadio, J.M., Towbin, A.J., Vaughan, M.R., Triplet, M., Ott-Napier, K., Dishman, D.J., Backus, L.R., Stockman, B., Brunner, M., Simpson, K., Spavin, R., Conner, J. and Cripe, T.P. 2017. Intratumoral injection of HSV1716, an oncolytic herpes virus, is safe and shows evidence of immune response and viral replication in young cancer patients. Clinical Cancer Research. 23(14), pp.3566–3574.

Tan, Z., Wu, Y., Fan, Z., Gao, T., Ding, S., Han, L., Luo, S., Fan, Q., Shi, J., Bai, C., Xue, R., Li, S., Zhang, L., Wang, X., Jia, L., Zhou, L., Liu, B., Huang, J. and Liu, J. 2025. Intratumoral oncolytic virus OH2 injection in patients with locally advanced or metastatic sarcoma: a phase 1/2 trial. Journal for immunotherapy of cancer. 13(1), e010543.

Tawbi, H.A., Burgess, M., Bolejack, V., Van Tine, B.A., Schuetze, S.M., Hu, J., D’Angelo, S., Attia, S., Riedel, R.F., Priebat, D.A., Movva, S., Davis, L.E., Okuno, S.H., Reed, D.R., Crowley, J., Butterfield, L.H., Salazar, R., Rodriguez-Canales, J., Lazar, A.J., Wistuba, I.I., Baker, L.H., Maki, R.G., Reinke, D. and Patel, S. 2017. Pembrolizumab in advanced soft-tissue sarcoma and bone sarcoma (SARC028): a multicentre, two-cohort, single-arm, open-label, phase 2 trial. The Lancet. 18(11), pp.1493–1501.

Taylor, A.M., Sheng, J., Ng, P.K.S., Harder, J.M., Kumar, P., Ahn, J.Y., Cao, Y., Dzis, A.M., Jillette, N.L., Goodspeed, A., Bodlak, A., Wu, Q., Isakoff, M.S., George, J., Grassmann, J.D.S., Luo, D., Flynn, W.F., Courtois, E.T., Robson, P., Hayashi, M., Paolillo, A.T., Petrilli, A.S., Caminada de Toledo, S.R., Balarezo, F.S., Lindsay, A.D., Hoang, B., Wong, S.T.C. and Lau, C.C. 2025. Immunosuppressive Tumor Microenvironment of Osteosarcoma. Cancers. 17(13), p.2117.

Tijtgat, J., De Munck, J., Dufait, I., Schwarze, J.K., Van Riet, I., Franceschini, L., Breckpot, K., Aerts, J.L., Neyns, B. and Tuyaerts, S. 2021. Unraveling the Effects of a Talimogene Laherparepvec (T-VEC)-Induced Tumor Oncolysate on Myeloid Dendritic Cells. Frontiers in immunology. 12, p.733506.

Todo, T., Ito, H., Ino, Y., Ohtsu, H., Ota, Y., Shibahara, J. and Tanaka, M. 2022. Intratumoral oncolytic herpes virus G47Δ for residual or recurrent glioblastoma: a phase 2 trial. Nature Medicine. 28(8), pp.1630–1639.

Todo, T., Martuza, R.L., Rabkin, S.D. and Johnson, P.A. 2001. Oncolytic herpes simplex virus vector with enhanced MHC class I presentation and tumor cell killing. Proceedings of the National Academy of Sciences of the United States of America. 98(11), pp.6396– 401.

Tu, B., Du, L., Fan, Q.-M., Tang, Z. and Tang, T.-T. 2012. STAT3 activation by IL-6 from mesenchymal stem cells promotes the proliferation and metastasis of osteosarcoma. Cancer letters. 325(1), pp.80–8.

Valyi-Nagy, T., Fareed, M.U., O’Keefe, J.S., Gesser, R.M., MacLean, A.R., Brown, S.M., Spivack, J.G. and Fraser, N.W. 1994. The herpes simplex virus type 1 strain 17+ gamma 34.5 deletion mutant 1716 is avirulent in SCID mice. The Journal of general virology. 75 ( **Pt 8**), pp.2059–63.

Voeller, J. and Sodel, P.M. 2019. Advances in Anti-GD2 Immunotherapy for Treatment of High Risk Neuroblastoma. Journal of Pediatric Hematology/Oncology. 41(3), pp.163– 169.

Wang, H., Borlongan, M., Kaufman, H.L., Le, U., Nauwynck, H.J., Rabkin, S.D. and Saha, D. 2024. Cytokine-armed oncolytic herpes simplex viruses: a game-changer in cancer immunotherapy? Journal for ImmunoTherapy of Cancer. 12(5), p.e008025.

Wang, S., Yan, W., Kong, L., Zuo, S., Wu, J., Zhu, C., Huang, H., He, B., Dong, J. and Wei, J. 2023. Oncolytic viruses engineered to enforce cholesterol efflux restore tumor-associated macrophage phagocytosis and anti-tumor immunity in glioblastoma. Nature Communications. 14(1), p.4367.

Wang, Y., Liu, X., Xie, G. and Li, Z. 2026. Immune Infiltration Landscape in Osteosarcoma: Clinical Implications for Prognosis and Therapy. Cancer Reports. 9(2): e70495

Wantoch, M., Wilson, E. B., Droop, A. P., Phillips, S. L., Coffey, M., El-Sherbiny, Y. M., Holmes, T. D., Melcher, A. A., Wetherill, L. F., & Cook, G. P. (2022). Oncolytic virus treatment differentially affects the CD56^dim^ and CD56^bright^ NK cell subsets in vivo and regulates a spectrum of human NK cell activity. Immunology, 166(1), 104–120.

Wedekind, M.F., Miller, K.E., Chen, C.-Y., Wang, P.-Y., Hutzen, B.J., Currier, M.A., Nartker, B., Roberts, R.D., Boon, L., Conner, J., LaHaye, S., Kelly, B.J., Gordon, D., White, P., Mardis, E.R. and Cripe, T.P. 2021. Endogenous retrovirus envelope as a tumor-associated immunotherapeutic target in murine osteosarcoma. iScience. 24(7), p.102759.

Whelan, J.S., Jinks, R.C., McTiernan, A., Sydes, M.R., Hook, J.M., Trani, L., Uscinska, B., Bramwell, V., Lewis, I.J., Nooij, M.A., van Glabbeke, M., Grimer, R.J., Hogendoorn, P.C.W., Taminiau, A.H.M. and Gelderblom, H. 2012. Survival from high-grade localised extremity osteosarcoma: combined results and prognostic factors from three European Osteosarcoma Intergroup randomised controlled trials. Annals of oncology : official journal of the European Society for Medical Oncology. 23(6), pp.1607–16.

Wilson, B.J., Owston, H.E., Iqbal, N., Giannoudis, P. V., McGonagle, D., Pandit, H., Philipose Pampadykandathil, L., Jones, E. and Ganguly, P. 2024. In Vitro Osteogenesis Study of Shell Nacre Cement with Older and Young Donor Bone Marrow Mesenchymal Stem/Stromal Cells. Bioengineering. 11(2), p.143.

Workenhe, S.T., Simmons, G., Pol, J.G., Lichty, B.D., Halford, W.P. and Mossman, K.L. 2014. Immunogenic HSV-mediated oncolysis shapes the antitumor immune response and contributes to therapeutic efficacy. Molecular therapy : the journal of the American Society of Gene Therapy. 22(1), pp.123–31.

Xie, L., Xu, J., Sun, X., Guo, W., Gu, J., Liu, K., Zheng, B., Ren, T., Huang, Y., Tang, X., Yan, T., Yang, R., Sun, K., Shen, D. and Li, Y. 2020. Apatinib plus camrelizumab (anti- PD1 therapy, SHR-1210) for advanced osteosarcoma (APFAO) progressing after chemotherapy: a single-arm, open-label, phase 2 trial. Journal for immunotherapy of cancer. **8**(1):e000798

Xu, W., Bian, Z., Fan, Q., Li, G. and Tang, T. 2009. Human mesenchymal stem cells (hMSCs) target osteosarcoma and promote its growth and pulmonary metastasis. Cancer Letters. 281(1), pp.32–41.

Yan, L., Guo, J., Zhong, Y., Wei, J. and Wang, Z. 2025. Molecular Mechanisms of Cell-to-Cell Transmission in Human Herpesviruses. Viruses. 17(6):742

Yu, A.L., Gilman, A.L., Ozkaynak, M.F., London, W.B., Kreissman, S.G., Chen, H.X., Smith, M., Anderson, B., Villablanca, J.G., Matthay, K.K., Shimada, H., Grupp, S.A., Seeger, R., Reynolds, C.P., Buxton, A., Reisfeld, R.A., Gillies, S.D., Cohn, S.L., Maris, J.M., Sondel, P.M. and Children’s Oncology Group 2010. Anti-GD2 antibody with GM-CSF, interleukin-2, and isotretinoin for neuroblastoma. The New England journal of medicine. 363(14), pp.1324–34.

Zheng, Y., Pei, Y., Dong, C., Liang, J., Cai, T., Zhang, Y., Tan, D., Wang, J. and He, Q. 2025. Oncolytic Herpes Simplex Virus Therapy: Latest Advances, Core Challenges, and Future Outlook. Vaccines. 13(8): 880.

Zhou, Z., Pang, Y., Ji, J., He, J., Liu, T., Ouyang, L., Zhang, W., Zhang, X.-L., Zhang, Z.-G., Zhang, K. and Sun, W. 2024. Harnessing 3D in vitro systems to model immune responses to solid tumours: a step towards improving and creating personalized immunotherapies. Nature Reviews Immunology. 24(1), pp.18–32.

