## Supplementary information for "Oncolytic virus-antibody combinations enhance immune-mediated killing of osteosarcoma"

Supplementary Figures, legends and tables:

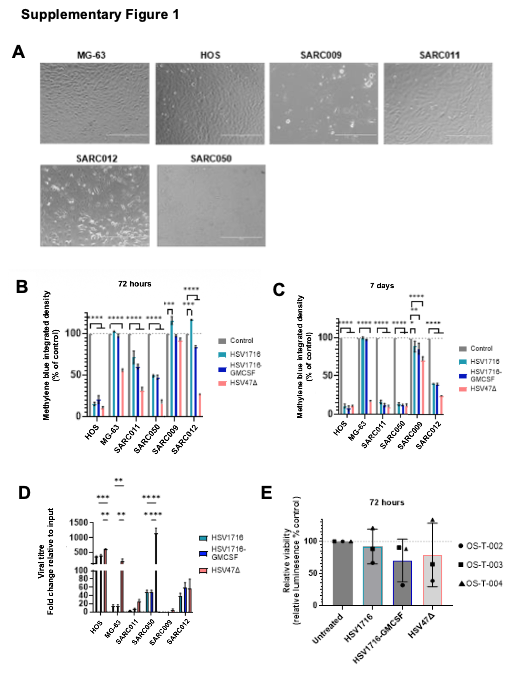

**Supplementary Figure 1: oHSVs exhibit direct oncolytic effects against OS cell lines and primary cell cultures.** (A) MG-63 and HOS OS cell lines and SARC011, SARC050, SARC009 and SARC012 primary cell cultures were seeded into 12 well plates at 1x10^5^ cells/well for 24 hours. Cells were then imaged using EVOS microscope at 10x magnification, scale bar = 400 µm. (B-C) OS cell lines and primary cell cultures were treated ± HSV1716, HSV1716-GMCSF or HSV47△ at 0.1 PFU/cell for (B) 72 hours or (C) 7 days. Monolayers were then stained with methylene blue, and the proportion of remaining viable cells was quantified by using ImageJ software to calculate integrated density of staining relative to untreated controls, mean ± SEM for n=3 independent experiments. Statistical comparison performed using two-way ANOVA. (D) OS cell lines and primary cell cultures were treated ± HSV1716, HSV1716-GMCSF or HSV47△at 1 PFU/cell for 72 hours, lysates were harvested and viral titre quantified using standard plaque assay on Vero cells. Results are presented as fold change in viral titre relative to input, mean ± SEM for n=3 independent experiments. Statistical comparison performed using two-way ANOVA. (E) OS patient tumours (OS-T-002, OS-T-003 and OS-T004) were digested in collagenase II and single cell suspensions were then added to low attachment 96 well plates. Cells were treated ± HSV1716-GFP, HSV1716-GMCSF or HSV47△ at 1 PFU/cell for 72 hours. Cell Titre-Glo® reagent was then added to wells and luminescence quantified. Results shown as percent luminescence relative to untreated controls, mean ± SEM for n=3 independent experiments.

**Supplementary Figure 2**

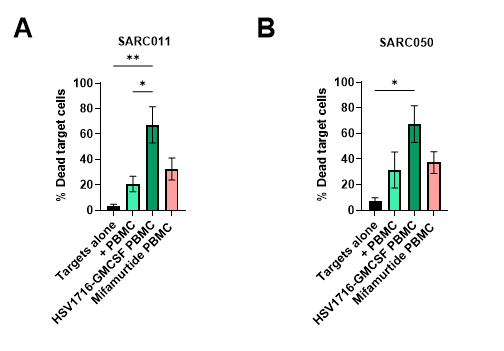

**Supplementary Figure 2: oHSVs stimulate immune-mediated killing of OS primary cell cultures**. (A-B) Healthy donor PBMC were activated ± HSV1716-GMCSF at 0.1 PFU/cell or Mifamurtide at 100 µM for 48 hours, PBMC were then co-cultured with CellTracker Green CMFDA stained SARC011 or SARC050 cells for 24 hours at a ratio of 20:1. Cells were then stained with yellow LIVE/DEAD stain, and the percentage of CellTracker Green + LIVE/DEAD+ target cells was quantified by flow cytometry. Results presented as mean ± SEM, n=4. Statistical comparison performed using one-way ANOVA.

**Supplementary Figure 3**

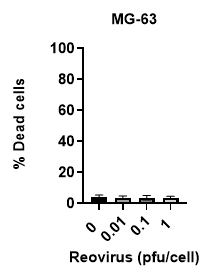

**Supplementary Figure 3: Oncolytic reovirus has no significant oncolytic effects against MG-63 OS cell line.** MG-63 cells were treated ± reovirus at 0.01, 0.1 or 1 PFU/cell for 72 hours. Cells were then stained with yellow LIVE/DEAD stain, and the percentage of dead cells quantified by flow cytometry. Results presented as mean ± SEM, n=3.

**Supplementary Table 1: Cell lines, primary cell cultures and patient sample information**

| **Cell type** | **Origin** | **Age** | **Sex** | **Prior treatment** | **Growth medium** |
| --- | --- | --- | --- | --- | --- |
| **Cell lines** | | | | | |
| MG-63 | Osteosarcoma, fibroblast-like | 14 | Male | Unknown | EMEM + 10% FBS |
| HOS | Osteosarcoma, epithelial-like with some fibroblast-like characteristics | 13 | Female | Unknown | EMEM + 10% FBS |
| Vero | African green monkey kidney epithelial cells | Unknown | Female | Not applicable | DMEM + 10% FBS |
| **Primary cell cultures** | | | | | |
| SARC011 | Osteosarcoma, chondroblastic | 12 | Female | Treatment naïve | DMEM + 10% FBS + 100 units of penicillin and 0.1mg/mL streptomycin |
| SARC050 | Osteosarcoma, osteoclast-rich, high grade | 28 | Female | Treatment naïve | DMEM + 10% FBS + 100 units of penicillin and 0.1mg/mL streptomycin |
| SARC009 | Osteosarcoma, osteoblastic central, high grade | 16 | Female | MAP chemotherapy | DMEM + 10% FBS + 100 units of penicillin and 0.1mg/mL streptomycin |
| SARC012 | Osteosarcoma, high-grade osteoblastic and chondroblastic | 17 | Male | MAP chemotherapy | DMEM + 10% FBS + 100 units of penicillin and 0.1mg/mL streptomycin |
| Mesenchymal stem cells(MSC) | Human Bone marrow-derived mesenchymal stem cells | 22 | Male | N/A | StemMACS™ MSC Expansion Media +100 units of penicillin and 0.1 mg/mL streptomycin |
| Peripheral blood mononuclear cells | Human healthy donor | Unknown | Unknown | N/A | RPMI + 10% FBS |
| **Patient tumour tissue samples** | | | | | |
| OS-T-002 | High grade osteosarcoma | 36 | Male | MAP chemotherapy | DMEM + 10 µg/ mL Fungin^TM^ + 10 µg/ mL Primocin^TM^ + 10% FBS |
| OS-T-003 | High grade osteoblastic osteosarcoma | 23 | Male | MAP chemotherapy | DMEM + 10 µg/ mL Fungin^TM^ + 10 µg/ mL Primocin^TM^ + 10% FBS |
| OS-T-004 | High grade osteosarcoma | 38 | Female | MAP chemotherapy | DMEM + 10 µg/ mL Fungin^TM^ + 10 µg/ mL Primocin^TM^ + 10% FBS |
| **Patient peripheral blood mononuclear cells** | | | | | |
| OS-B-002 | Osteosarcoma patient peripheral blood sample collected at point of surgery | 36 | Male | MAP chemotherapy | RPMI + 10% FBS |
| OS-B-003 | Osteosarcoma patient peripheral blood sample collected at point of surgery | 23 | Male | MAP chemotherapy | RPMI + 10% FBS |
| OS-B-004 | Osteosarcoma patient peripheral blood sample collected at point of surgery | 38 | Female | MAP chemotherapy | RPMI + 10% FBS |
| **Luc2-expressing cell cultures** | | | | | |
| **Cell type** | **Concentration of puromycin for selection (µg/mL)** | | | | |
| MG-63-Luc2 | 4 | | | | |
| HOS-Luc2 | 4 | | | | |
| SARC011-Luc2 | 2 | | | | |
| SARC050-Luc2 | 2 | | | | |

RPMI; Roswell Park Memorial Institute, DMEM; Dulbecco’s Modified Eagles Medium, EMEM; Eagles Minimum Essential Medium

**Supplementary Table 2: Antibodies, fluorescent stains and other reagents**

| **Flow cytometry antibodies** | | | | | | |
| --- | --- | --- | --- | --- | --- | --- |
| **Target protein** | **Target species** | **Isotype** | | | **Fluorophore** | **Manufacturer** |
| CD3 | Human | Mouse IgG2ak | | | PE | BD Biosciences |
| CD56 | Human | Mouse IgG2bk | | | BV421 | BD Biosciences |
| CD69 | Human | Mouse  IgG1k | | | BV605 | BD Biosciences |
| CD107a | Human | Mouse IgG1k | | | BV605 | BD Biosciences |
| CD14 | Human | Mouse IgG2bk | | | BUV395 | BD Biosciences |
| CD163 | Human | Mouse IgG1k | | | PE-CF594 | BD Biosciences |
| CD206 | Human | Mouse IgG1k | | | BV421 | BD Biosciences |
| HLA-DR  (MHC class II) | Human | Mouse IgG2ak | | | APC | BD Biosciences |
| EGFR | Human | Mouse IgG1κ | | | BB700 | BD Biosciences |
| GD2 | Human | Mouse IgG2a | | | BUV395 | BD Biosciences |
| CD340 (HER2) | Human | Mouse IgG1κ | | | BV421 | BD Biosciences |
| CD20 | Human | Mouse IgG2bk | | | APC | BD Biosciences |
| **Monoclonal antibodies** | | | | | | |
| **Antibody** | **Target protein** | **Concentration for cell treatments (µg/mL)** | | **Manufacturer** | | |
| Imgatuzumab | EGFR | 10 | | MedChem Express | | |
| Naxitamab | GD2 | 10 | | MedChem Express | | |
| Trastuzumab | HER2 | 10 | | TargetMol Chemicals Inc | | |
| Rituximab | CD20 | 10 | | MedChem Express | | |
| **ELISA antibodies** | | | | | | |
| **Product** | | | **Manufacturer** | | | |
| IFN-α mAb (MT1/3/5), unconjugated | | | Mabtech | | | |
| IFN-α mAb (MT2/4/6), biotin | | | Mabtech | | | |
| IL-6 Monoclonal Antibody (MQ2-13A5) | | | Invitrogen | | | |
| IL-6 Monoclonal Antibody (MQ2-39C3) biotin | | | Invitrogen | | | |
| GM-CSF mAb (21C11), unconjugated | | | Mabtech | | | |
| GM-CSF mAb (23B6), biotin | | | Mabtech | | | |
| Human IL-1 beta/IL-1F2  Quantikine ELISA Kit | | | Bio-techne | | | |
| **Other reagents** | | | | | | |
| **Product** | | | **Manufacturer** | | | |
| CellTiter-Glo® reagent | | | Promega | | | |
| Luciferin Potassium Salt | | | REGIS Technologies | | | |
| CellTracker Green CMFDA | | | ThermoFisher Scientific | | | |
| LIVE/DEAD™ Fixable Yellow Dead Cell Stain Kit | | | ThermoFisher Scientific | | | |
| Lympholyte^TM^ | | | Cedar Lane | | | |
| Brefeldin A Solution (1000X) | | | ThermoFisher | | | |
| Hanks Balanced Salt solution | | | Sigma | | | |
| Foetal Bovine Serum | | | Sigma | | | |
| CD14 Microbeads Human | | | Miltenyi Biotec | | | |
| Fungin | | | InvivoGen | | | |
| Primocin | | | InvivoGen | | | |
| Collagenase II | | | Invitrogen | | | |
| Mifamurtide | | | Cambridge Bioscience | | | |
| **Buffers** | | | | | | |
| MACS buffer | | | PBS + 1% FBS + 0.4% 0.5M EDTA | | | |
| FACS buffer | | | PBS + 10% FBS + 0.1% sodium azide | | | |
